# Exploring the fog vegetation in the marginal Atacama Desert: common evolutionary history of species-rich genera and morphological convergence in *Nolana* and *Heliotropium* sect. *Cochranea*

**DOI:** 10.1101/2025.09.04.673165

**Authors:** Sergio Castro

## Abstract

The flora of the Atacama–Peru Desert (7°–30°S) is characterized by its remarkable diversity and adaptation in the driest place on Earth. It is composed of endemic and diverse genera, with most species having restricted geographic distributions. To infer its evolutionary history, the structure of a fog-dependent plant community in the Marginal Desert (27°–30°S) was characterized, with a special focus on congeners of *Nolana* and *Heliotropium* sect. *Cochranea*. Similarity profile analysis (SIMPROF) identified six plant assemblages differentially distributed along the altitudinal gradient. The congeners showed a pattern of niche segregation, with basal species *N. rostrata* and *H. filifolium* occupying the fog-influenced altitudinal range, while recently diverged species occurred outside its direct influence. To corroborate phylogenetic convergence, climatic zones were defined based on the presence of *Nolana* and *Cochranea*, and these were used to reconstruct ancestral areas and formulate a biogeographic hypothesis. The history of the Atacama Desert flora is marked by two periods: the formation of the hyperarid core (18°–27°S) during the late Miocene led to an initial divergence, leaving species isolated in different climatic zones. Then, beginning around 4 Mya, various wet phases promoted the northward geographic expansion of the flora inhabiting the Marginal Desert, with some genera reaching as far as Peru. Following geographic expansion, dry phases led to landscape-scale heterogeneity in water availability, promoting adaptive radiation. The ecological and phylogenetic affinities of *N. rostrata* and *H. filifolium* suggest that terete leaf morphology represents a case of evolutionary convergence associated with fog water capture.

## 1. Introduction

Arid and semi-arid regions are characterized by water scarcity and by hosting habitats with highly variable water availability, temporally and spatially (Stebbins, 1952; Gaddis et al., 2016). The rapid origin, climax, and decline of plant species, which is a hallmark of arid and semi-arid regions (Stebbins, 1952), results in original diversity of genera and species (Dillon et al., 2009; Folk et al., 2024). Given that plants are immobile, they are highly susceptible to any lasting change in their habitat. As a result, plants that have evolved in arid environments exhibit a wide range of adaptations that enhance the acquisition (Mooney et al., 1980; Rundel et al., 1980), optimization (Mazer et al., 2020), and storage of their resources (Tribble et al., 2021). Unique evolutionary innovations have arisen as a result of plant adaptation to water scarcity, such as the succulent growth form (Guerrero et al., 2019) or CAM photosynthesis (Chomentowska et al., 2025). Thus, closely related taxa that make up xerophytic lineages are valuable examples for the study of evolution, as they allow for the placement of morphological adaptations within a phylogenetic framework and the subsequent formulation of hypotheses about the causes and consequences of lineage diversification (Castro et al., 2023).

The Atacama-Peru Desert (7°–30°S) is located on the western coast of South America and spans 3,500 km in length and 80–160 km in width. It is characterized by its biodiversity, which is largely isolated from neighboring regions (Rundel et al., 1991; Gengler–Nowak, 2002), and by being the driest desert on Earth, with some areas averaging less than 1 mm of annual precipitation (Trewartha, 1961; Rundel et al., 1991; Schulz et al., 2012). To understand the causes of the region’s extreme aridity, it is necessary to consider its geography. In the Chilean portion of the desert, a hyper-arid core develops northward from the 27°S parallel between the Pacific Ocean and the Andes Mountains (Fig. 1) (Houston & Hartley, 2003; Dunai et al., 2005). To the west of this core, the incursion of moist air from the Pacific Ocean is blocked by a subsidence inversion, a meteorological phenomenon caused by the South Pacific Anticyclone and intensified by the Humboldt Cold Current, which significantly limits the formation of high-altitude clouds (cirrocumulus), a source of precipitation (Gengler–Nowak, 2002; Larrain et al., 2002; Schulz et al., 2012). To the east, a classic rain shadow effect produced by the Andes traps moisture coming from the Amazon basin and the Atlantic Ocean (Houston & Hartley, 2003). These meteorological phenomena driven by the interaction between large-scale movements of water and air with specific geographic conditions result in a precipitation gradient that decreases from south to north (Houston & Hartley, 2003; Schulz et al., 2012).

**Figure 1.**
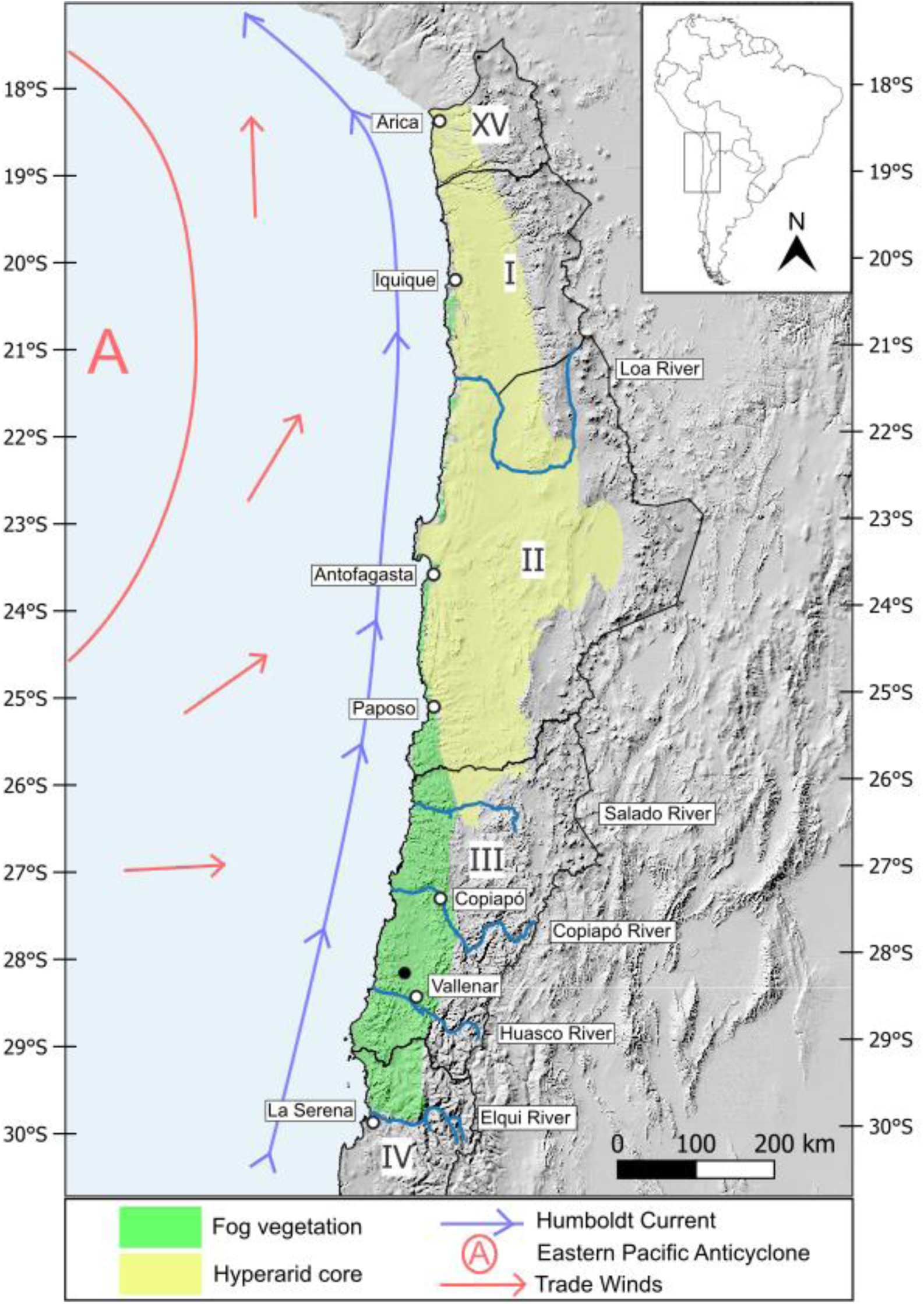
Chilean portion of the Atacama – Peru Desert. Open dots are cities and villages. The black dot represents the study site. Roman numbers represent the numeration of Chilean political reg

As a result, the moisture necessary to sustain most of plant life is only available in the areas surrounding the hyper-arid core: on the eastern side, at the foothills of the Andes, plants grow thanks to watersheds originating in the Andean peaks that flow westward during the Altiplanic winter; on the western coastal strip, the main source of moisture comes from large fog banks that penetrate several kilometers inland, depending on atmospheric and orographic factors (Cereceda et al., 2002). These fog banks are generally of advective origin, with low clouds (stratocumulus) forming over the ocean below the thermal inversion layer, between 400 and 1,500 meters above sea level (Espejo, 2001), and being transported to the coast by trade winds. Orographic fog also forms, more locally, in areas with particular conditions such as hilltops surrounded by coastal zones with upwelling of cold water (Cereceda et al., 2002). When advective or orographic fog collides with coastal hills that rise above the thermal inversion layer, it generates underground water flows that allow for plant growth (Rundel et al., 1991). These flows can even be seen discharging into large open-pit mines. The dynamics and behavior of fog banks remain not fully understood (Gonzales et al., 2023), but locations hundreds of kilometers apart have been reported to exhibit the same patterns of fog abundance variation throughout the year (Osses et al., 1998; Larrain et al., 2002), and the influence of the El Niño Southern Oscillation (ENSO) has also been suggested (Muñoz-Schick et al., 2001; Eichler & Londoño, 2013).

The contribution of moisture from fog becomes essential at a certain point along the precipitation gradient. As this occurs, fog-dependent vegetation shifts from covering several kilometers inland in the southern portion of the Atacama Desert (i.e., the “Marginal Desert”, 27°–30°S) to becoming restricted only to the slopes of hills adjacent to the ocean (Fig. 1). In the locality of Paposo (25°S), for example, vegetation forms a “fertile belt” that grows within the fog’s altitudinal range (300–800 meters above sea level) (Johnston, 1929), and the transition between shrubland with 50% plant cover and absolute desert occurs over just a few hundred meters (Rundel & Mahu, 1976). Farther north, fog vegetation becomes very sparse, appearing as isolated patches on limited portions of the hillsides (Muñoz-Schick et al., 2001; Luebert et al., 2007; Schulz et al., 2011) and eventually disappears altogether, creating a notable biogeographic gap at the Chile–Peru border (18°S) (Dillon et al., 2009; Rundel et al., 1991; Pinto & Luebert, 2009; Luebert, 2011a; Ruhm et al., 2022). The taxonomic composition of coastal vegetation changes from south to north (Schulz et al., 2011), while species richness appears to decline though not functional diversity (Stotz et al., 2021).

The boundaries of the Atacama Desert have historically been a subject of debate (Luebert, 2011). However, the Elqui River, located at the 30°S parallel (Fig. 1), is considered the boundary between the semi-arid and Mediterranean regions in Chile (Etienne & Prado, 1982); it marks the distribution limit of several species typical of the Mediterranean sclerophyllous forest and signals the beginning of what some authors have called the Marginal Desert, which extends northward to the 27°S parallel (Gengler–Nowak, 2002). This biogeographic region largely corresponds to the Chilean political region known as the Atacama Region or Third Region (“III Region” hereafter; 26°–29°20′S), and is notable for the diversity and endemism patterns of its flora. In the Marginal Desert, there are peaks in the diversity of endemic and native Chilean cacti (Guerrero et al., 2011). Additionally, the vascular flora of the III Region contains 19.3% (1,099 species) of the total plant species present in continental Chile (Letelier et al., 2008). Of these, 54.3% are endemic to the country, 37.3% are endemic to Regions II, III, and IV (Fig. 1), and 7.9% are endemic solely to the III Region, with the shrub growth form having the highest number of endemic taxa. Notably, 54% of Chile’s native flora and 64% of its endemic flora reach either their northern or southern distribution limits within the III Region (Squeo et al., 2008).

In the Marginal Desert and the coastal strip north of the 27°S parallel, there are endemic plant lineages of Chile and Peru that diversified in situ and include a considerable number of species: *Nolana* (Solanaceae, 92 spp.) (Dillon, 2023), *Heliotropium* sect. *Cochranea* (“*Cochranea*” hereafter, Boraginaceae, 17 spp.) (Luebert, 2013), *Copiapoa* (Cactaceae, 32 spp.) (Larridon et al., 2015), *Cristaria* (Malvaceae, ∼20 spp.) (Muñoz-Schick, 1995), and *Eulychnia* (Cactaceae, ∼8–10 spp.) (Larridon et al., 2018; Merklinger et al., 2021). It is common to see several species from these genera growing in sympatry, just a few meters apart (Dillon et al., 2009; Luebert et al., 2014), and it can be striking in the field to observe such marked morphological differences among species that appear to share highly overlapping niches. This raises questions such as: What conditions drove the diversification of these lineages, resulting in species with such distinct morphologies yet seemingly similar ecological requirements? Or how did these lineages diversify if their species do not appear to have undergone ecological differentiation? The most commonly proposed diversification models follow the pattern of allopatric speciation, based on the idea that climatic changes such as alternating pluvial phases and prolonged arid periods, along with glacial cycles led to isolation among plant populations in the Atacama Desert, which promoted speciation through genetic differentiation (Luebert & Wen, 2008; Dillon et al., 2009; Böhnert et al., 2022; Merklinger et al., 2021; Meriño et al., 2024). It has also been suggested that the rapid uplift of the Andes during the late Neogene created the current hyper-arid conditions (Houston & Hartley, 2003; Luebert et al., 2011b), promoting habitat fragmentation (Merklinger et al., 2021). In contrast, few studies have discussed the occurrence of local ecological speciation events in sympatry.

This study aims to contribute to our understanding of plant speciation events and their history in the Atacama Desert. Preliminary observations from a vegetation survey in the Marginal Desert suggest spatial segregation between *Heliotropium filifolium* and *H. myosotifolium* (Fig. S2). Additionally, four species of the genus *Nolana* and three of *Cochranea* inhabit the study site. Therefore, considering the region’s high endemism, particularly among shrub life forms, the composition and local distribution of plant assemblages at the study site can provide insights into habitat differences that have played a role in the speciation processes responsible for the current diversity and ecological patterns. As such, treating plant assemblages as units that host ecologically similar species, this study aimed to: (i) delimit the plant assemblages in the study site based on species composition and abundance, and to (ii) determine the effects of elevation, slope, and aspect on their distribution. During the course of the analyses, it was observed that the species of *Nolana*, *Cochranea*, and *Copiapoa* occurring within the fog altitudinal range belonged to basal clades in their respective phylogenies. Therefore, to explore a possible case of phylogenetic convergence, (iii) climatic zones were defined within the Chilean portion of the Atacama Desert based on the distribution of *Nolana* and *Cochranea*, and these zones were used to (iv) reconstruct the biogeographic history of *Cochranea* in order to compare it with that of other genera in Atacama.

## 2. Materials and Methods

### 2.1 Study site and data collection

The study site is located in the Coastal Cordillera of Chile’s III Region, 30 km from the city of Vallenar and 36 km from the Pacific Ocean, in the vicinity of the “Los Colorados” iron mine operated by Compañía Minera del Pacifico (28°17′S) (Figs. 1, S1). The vegetation consists of low shrublands with less than 25% cover, succulent formations that can reach more than 75% cover, and annual herbs formations. According to the vegetation and climate classification by Luebert and Pliscoff (2006), the study area lies within the vegetation belts “Matorral desértico mediterráneo interior de *Skytanthus acutus* – *Atriplex deserticola*” and “Matorral desértico interior de *Adesmia argentea* – *Bulnesia chilensis*”, both characterized by a desert-oceanic Mediterranean bioclimate with low annual precipitation and low continentality.

To characterize the plant assemblages at the study site, sampling points were first established within independent plant formations based on satellite imagery. At each sampling point, 500 m² plots (50 x 10 meters) were created using measuring tapes, and the abundance of the species falling in was quantified. Samples were collected for subsequent laboratory identification of species that could not be identified in the field, based on available literature regarding distribution and morphology (Squeo et al., 2008; Rodríguez et al., 2018). Annual species and geophytes lacking structures necessary for taxonomic identification, or species that were identified but whose abundance was not precisely quantified, were excluded from subsequent analyses. Four field campaigns were conducted during March, May, August, and September 2023.

### 2.2 Community Analyses

To group the plant assemblages based on differences in species abundance and composition without any prior assumptions, a Similarity Profile Analysis (SIMPROF) was performed. This permutation-based technique assigns a significance value to the nodes of a clustering dendrogram, which in our analysis was constructed using the UPGMA method (“unweighted pair group method with arithmetic mean”) based on a Bray-Curtis dissimilarity matrix. Non-metric multidimensional scaling (nMDS) analysis was carried out to visually represent the taxonomic differences among the groups defined by the SIMPROF analysis. Indicator species for each vegetation formation group were identified using a multi-level pattern analysis, where a specificity or predictive value (value “A”) and a fidelity or sensitivity value (value “B”) are assigned to indicator species. Value A represents the probability that if a species is found in a formation, it corresponds to the target formation, and value B represents the probability of finding the indicator species within the target formation. Community indices (total abundance, species richness, Shannon diversity, and Pielou’s evenness) were calculated for each sampling plot and compared among Simprof groups using one-way analysis of variance (ANOVA) and Tukey’s post hoc test. When ANOVA residuals did not meet normality assumptions, the non-parametric Kruskal-Wallis test and Wilcoxon significance test were applied. All analyses were performed using the R packages vegan (Oksanen et al., 2016), FactoMineR (Lê et al., 2008) and dplyr (Yarberry, 2021).

Next, to determine the distribution of each group of plant assemblages with respect to the variables elevation, slope, and orientation (aspect), a high-resolution digital elevation model (DEM) was downloaded from the United States Geological Survey (USGS) website. Topographic variable values were extracted for each sampling plot using QGIS version 3.36.0. These values were analyzed using ANOVA and Tukey’s test. The aspect variable was transformed into a qualitative variable and a Correspondence Analysis (CA) was performed. For the *Nolana* and *Cochranea* species present at the study site, linear regressions were used to evaluate whether there was a relationship between their abundance and altitude, with ade4 package (Dray & Dufour, 2007),

### 2.3 Climatic Niche Analysis

With the aim of identifying significant climatic zones for the biogeographic reconstruction of the Atacama Desert genera, presence data for 15 species of *Nolana* and 10 species of *Cochranea* were downloaded from the Global Biodiversity Information Facility (GBIF, http://www.gbif.org/) and filtered based on the following criteria: 1. Human identification or specimen preserved in a herbarium; 2. Geographic precision less than 500 m; 3. Collection or sighting year more recent than 1950; 4. Consistency with the known geographic distribution of the species (Rodríguez et al., 2018). Considering the spatial and ecological segregation of *Heliotropium filifolium* and *H. myosotifolium* at the study site, vector layers of the 17 most informative climatic variables for these species (Pliscoff et al., 2014) were downloaded, based on the assumption that these variables would also be informative to distinguish the climatic niches of the other species in the analysis.

Climatic values for each presence record were subjected to Principal Component Analysis (PCA) and Discriminant Analysis (DA). In both cases, the objective was to visualize how and to what extent species cluster in multivariate space; however, the grouping method differs between analyses: in PCA, the creation of axes seeks to maximize total variation by grouping variables with the greatest variance, so points are distributed based on their overall similarity. In DA, the axes are created to group points according to a prior assignment to species, and thus these axes do not represent the greatest variance in the dataset but the variables that best discriminate between species. The accuracy of the DA model and the kappa coefficient were calculated using a confusion matrix. The R packages vegan (Oksanen et al., 2016), ade4, FactorMineR (Lê et al., 2008), factoextra (Kassambara & Mundt, 2017), caret (Kuhn, 2008), and lmtest (Hothorn et al., 2015) were used for analyses, and climatic variable values were extracted using QGIS version 3.36.0.

### 2.4 Phylogenetic Analyses and Ancestral Area Reconstruction

Sequences corresponding to one nuclear ribosomal locus (ITS) and four chloroplast loci (*ndhF*, *rps16*, *trnL-trnF*, and *trnS-trnG*) from 16 *Cochranea* species and 2 outgroups were compiled (Table S2). For the phylogenetic analysis, the five loci were aligned and concatenated using Geneious software (Kearse et al., 2012). Phylogenetic analyses were performed using Bayesian inference (BI) in MrBayes 3.2.7 (Ronquist et al., 2012), and the best nucleotide substitution model (GTR+I+G) was selected with jModeltest 2.1.3 (Darriba et al., 2012). The analysis consisted of two independent Markov Chain Monte Carlo (MCMC) runs with 10 million generations, sampling every 1000 generations after a burn-in period of 25% of the initial generations. Chain stationarity was confirmed with Tracer 1.7.1 (Rambaut et al., 2018), and the resulting tree was edited with FigTree v1.4.4 (Rambaut, 2019).

Ancestral areas were estimated using the S-DIVA analysis implemented in the RASP (Reconstruct Ancestral State Phylogenies) program v2.1 (Yu et al., 2015). The distribution ranges of the 16 *Cochranea* species included in the phylogenetic analysis were categorized within the previously defined climatic zones: A: North Coast, B: North Interior, C: South Coast, and D: South Interior. The S-DIVA analysis was run using the t. file obtained from the phylogenetic analysis, allowing a maximum of two ancestral areas per node in the tree.

## 3. Results

### 3.1 Floristic diversity

The abundance of 46 vascular plant species was quantified, comprising 7,252 individuals distributed across 144 plots, equivalent to 72,000 m² (7.2 ha) (Fig. S1). The sampling elevation range spanned 441 meters (from 356 to 797 m.a.s.l.). Among taxa that could not be quantified in terms of abundance, 19 were classified at the species level and 13 at the genus level, totaling 78 species present in the study area, belonging to 38 families (Table S1). The percentage of endemic species to Chile was 52.56%, native species 25.64%, and introduced species 5.13%.

The families with the greatest species richness were Asteraceae (10 species), Solanaceae (8 species), Fabaceae (7 species), Boraginaceae (6 species), and Cactaceae and Aizoaceae (4 species each). Two fern species and one gymnosperm were identified. The five most abundant species were *Encelia canescens* (Asteraceae), *Cumulopuntia sphaerica* (Cactaceae), *Eulychnia acida* (Cactaceae), *Miqueliopuntia miquelii* (Cactaceae), and *Heliotropium filifolium* (Boraginaceae) (Table S1).

### 3.2 Community structure

The SIMPROF analysis identified ten plant assemblages groups based on species composition and abundance (Fig. 2a), of which six were considered valid (Groups 3, 4, 5-6-7, 8, 9, and 10). Regarding topographic variables, orientation was the only one that did not show significant differences among the groups. In general, vegetation groups segregated along elevational ranges (Figs. 3, S3). The nMDS analysis revealed niche segregation of *Nolana* and *Cochranea* species (Fig. 2b). Linear models indicated that elevation significantly influenced the abundance *N. rostrata*, *H. filifolium*, and *N. divaricata* (Fig. S6).

**Figure 2.**
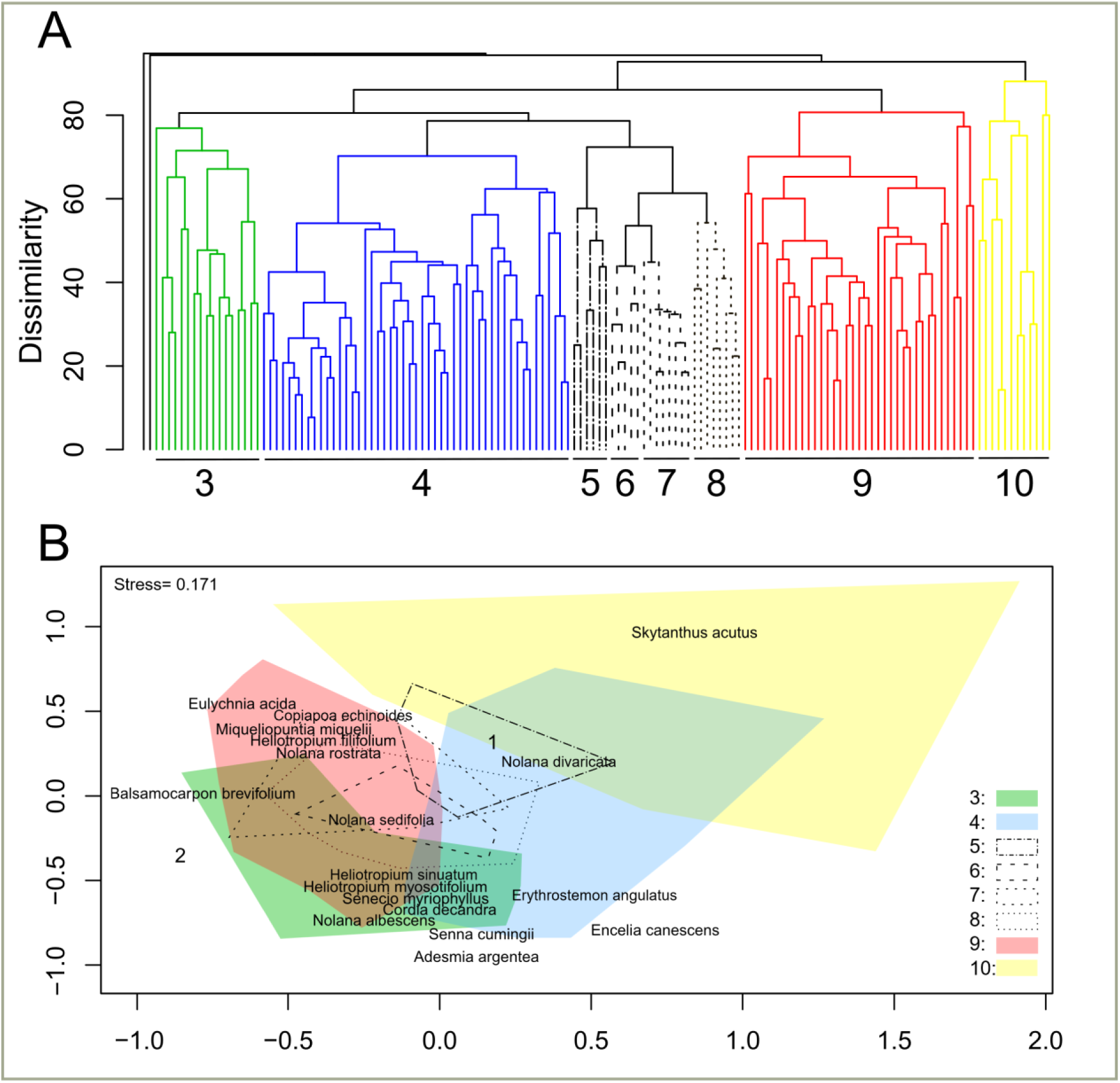
Community structure of fog vegetation in the study site. A, Similarity profile analysis (SIMPROF) based on a Bray-Curtis dissimilarity matrix. Colored and dashed clades represent clades that differ from each other on p < 0.05, thus defined as groups (types) of plant assemblages. Number of permutations= 999. B, nMDS analysis based on 999 permutations. Colored areas correspond to significant clades from SIMPROF analysis (see Fig S3 for point clouds). Species names correspond to species with high and significant R2 values (Appendix S1). Numbers correspond to plots that did not group with other assemblages in SIMPROF analysi

**Figure 3.**
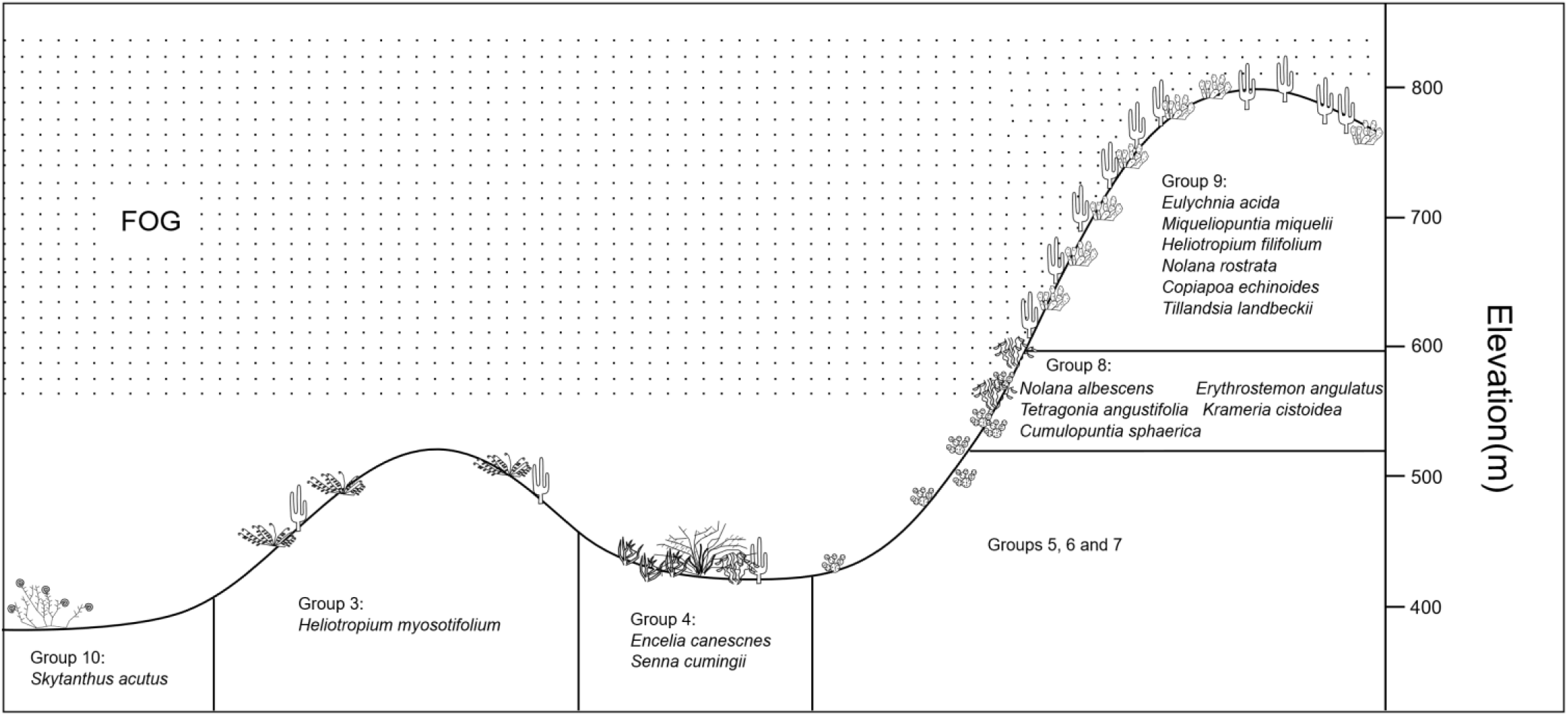
Diagramatic view of the fog vegetation at the study site, Los Colorados, Atacama, Chile. Indicator species of each group are shown. The fog position is hypothetical.

Groups 1 and 2 each included only one sampling plot and were not clustered with other plots due to their uniqueness: the plot from Group 1 included only three individuals of *Cumulopuntia sphaerica*, while the plot from Group 2, although containing six species, was dominated by *Balbisia peduncularis* (100 individuals).

Group 3 was sampled in 17 plots and corresponds to a low shrubland. It is characterized by the presence of *Heliotropium myosotifolium*, identified as the only indicator species (Appendix S2). Associated species included *Encelia canescens* and *Adesmia argentea*. This formation is distributed in the lower to mid-elevation range, from 397 to 515 m.a.s.l., on slopes between 10% and 20% (Fig. 3, S4).

Group 4 was the most represented in the sampling with 49 plots. The vegetation corresponds to low shrublands and occasionally to forests due to the presence of the tree species *Cordia decandra*. Its species composition is similar to Group 3, but with higher abundance of *Encelia canescens* and slightly greater species richness and total abundance, although not significantly different (Fig. S5). It is found in the low-elevation zone of the study area, in dry washes with slopes below 10%, distinguishing it from Group 3 which occurs on slopes (Figs. 3, S4). Indicator species are *Encelia canescens* and *Senna cumingii*.

Groups 5, 6, and 7 are distributed across the lower and mid-elevation zones (Fig. 3), have low plant cover, and lack clear indicator species. However, the most abundant species is *C. sphaerica*, whose abundance increases with elevation (Table S1). Thus, these three groups may represent a single vegetation type subdivided according to *C. sphaerica* abundance levels.

Group 8 corresponds to low shrublands. This type of formation is found within a limited range in the mid-to-upper sampling zone (487–597 m.a.s.l., with one outlier at 377 m a.s.l.) and shows the highest species richness and total abundance (Figs. S4, S5). Indicator species are *C. sphaerica*, *Tetragonia angustifolia*, *Krameria cistoidea*, *Nolana albescens*, and *Erythrostemon angulatus*, although all occur in other groups as well, resulting in low specificity values (A < 0.5; Appendix S2). It is the only group with a tendency toward a specific orientation (North-facing slopes) (Fig. S4).

Group 9 occurs at higher elevations, directly influenced by fog, between 426 and 797 m a.s.l., and on slopes steeper than 15%. It comprises shrublands with succulents or pure succulent formations. Its diversity, richness, and abundance values are high and similar to those of Group 4. Indicator species include the succulents *Eulychnia acida*, *Miqueliopuntia miquelii*, *Copiapoa echinoides*, and *Tillandsia landbeckii*, as well as the terete-leaved shrubs *Heliotropium filifolium* and *Nolana rostrata*. Notably, *N. rostrata* and *H. filifolium* share similar leaf morphology (Fig. 6), and along with *C. echinoides*, are located in similar clades within the phylogenies of their respective genera (Fig. 5; see section 4.2).

Finally, Group 10 includes plant assemblages in the lower sampling area, in flat zones with sparse shrub cover, corresponding to annual herbs formations with scattered shrubs. This group has the lowest richness and total abundance, with only one indicator species, *Skytanthus acutus*.

### 3.3 Climatic niche analysis

In the combined climatic niche analysis of *Nolana* and *Cochranea* species, the species clustered into four climatic zones in the multivariate PCA space (Figs. 4, S7). The first PCA axis explained 49.8% of the total variation and was correlated with temperature variables (Tmean1, r = 0.89; Tmin10, r = 0.84; Bio11, r = 0.82), and to a lesser extent with precipitation variables (Prec3, r = 0.39; Prec7, r = 0.32). This axis separated species distributed along the coastal strip north of 27°S (“Northern Coast”) from those found to the south, both on the coast (“Southern Coast”) and inland (“Southern Interior”), while species occurring in the hyperarid core north of 27°S (“Northern Interior”) clustered at the most negative values of the axis (Fig. 4a). The second axis was related to thermal amplitude variables (Bio2, r = 0.98; Bio3, r = 0.82) and separated species from the Northern and Southern Interior groups from those in the Northern and Southern Coast (Fig. 4a).

**Figure 4.**
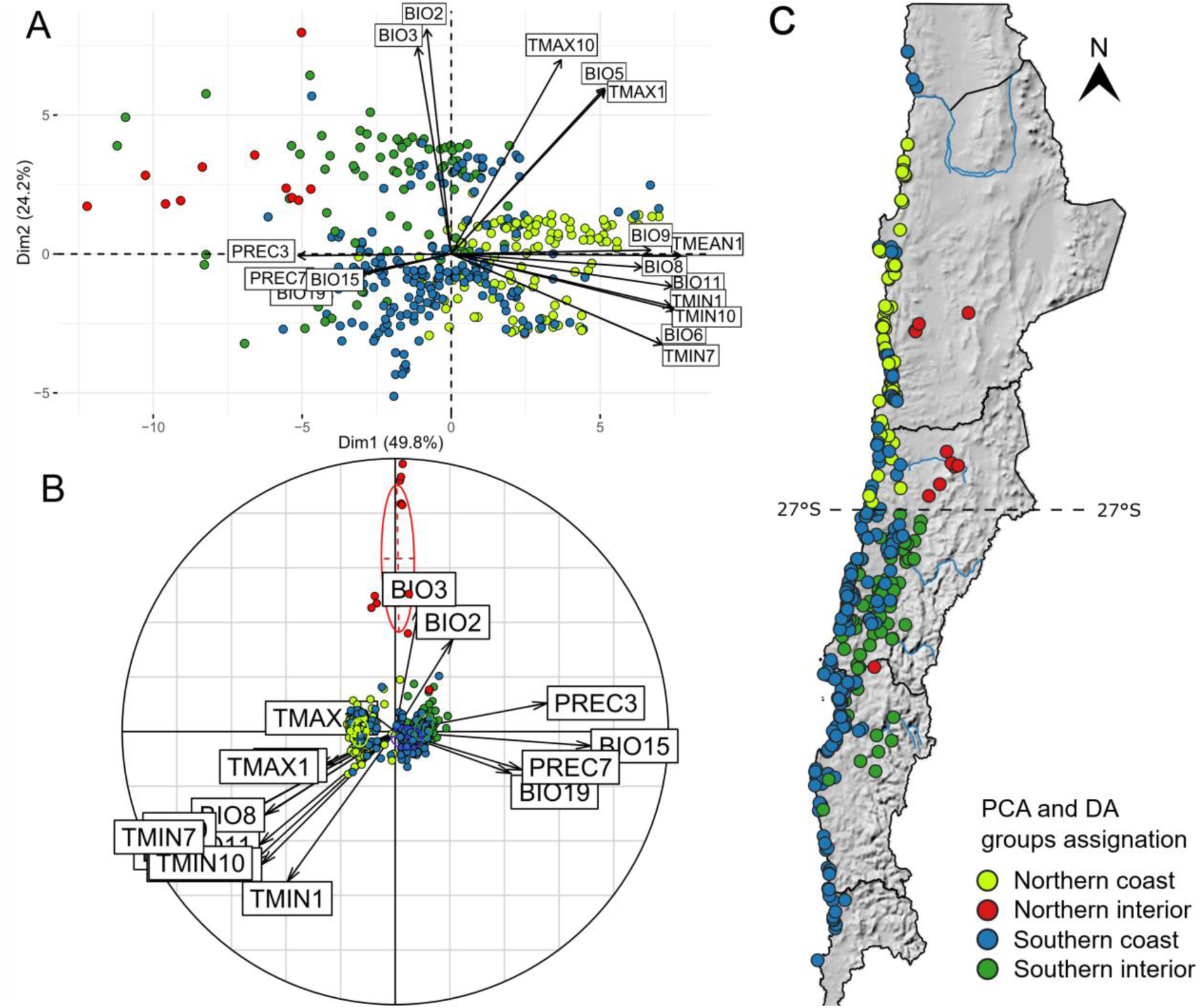
Climatic zones in the Atacama Desert based on the presence of *Nolana* and *Cochranea*. A, Principal component analysis of the climatic niche of 15 *Nolana* species and 10 *Cochranea* species. B, Discriminant analysis determining the best explicative variables for species climatic niche distinction (model accuracy: 0.85; қ coefficient: 0.75). See Fig S7 for species names in the multivariate space of both analyses and Fig S11 for single species presence data. C. Geographic distribution of data and corresponding climatic zone assignation. BIO2, Mean diurnal range; BIO3, Isothermality; BIO5, Maximal temperature of the warmest month; BIO6; Minimal temperature of coldest month; BIO8, Mean temperature of the wettest quarter; BIO9, Mean temperature of the driest quarter; BIO11, Mean temperature of the coldest quarter; BIO15, Precipitation seasonality; BIO19, Precipitation of the coldest quarter; PREC3, March precipitation; PREC7, July precipitation, TMAX1, January mean maximum temperature; TMAX10, October mean maximum temperature; TMEAN1, January mean temperature; TMIN1, January mean minimum temperature; TMIN7, July mean minimum temperature; TMIN10, October mean minimum temperature.

The Discriminant Analysis (DA) did not clearly differentiate between the Southern Coast and Southern Interior groups, but it distinguished species in the Northern Interior group, which occupy a climatic niche with a greater temperature range (Fig. 4b). On the other hand, the Northern Coast group was characterized by the highest temperatures, while the Southern Interior and Southern Coast groups had higher precipitation values (Fig. 5). When the species were grouped into four climatic zones as suggested by the PCA, the DA model had an accuracy of 0.83 and a Kappa coefficient of 0.74. However, when only three groups were considered (merging Southern Coast and Southern Interior), both accuracy and Kappa increased: accuracy = 0.93, Kappa = 0.86 (Appendix S3).

**Figure 5.**
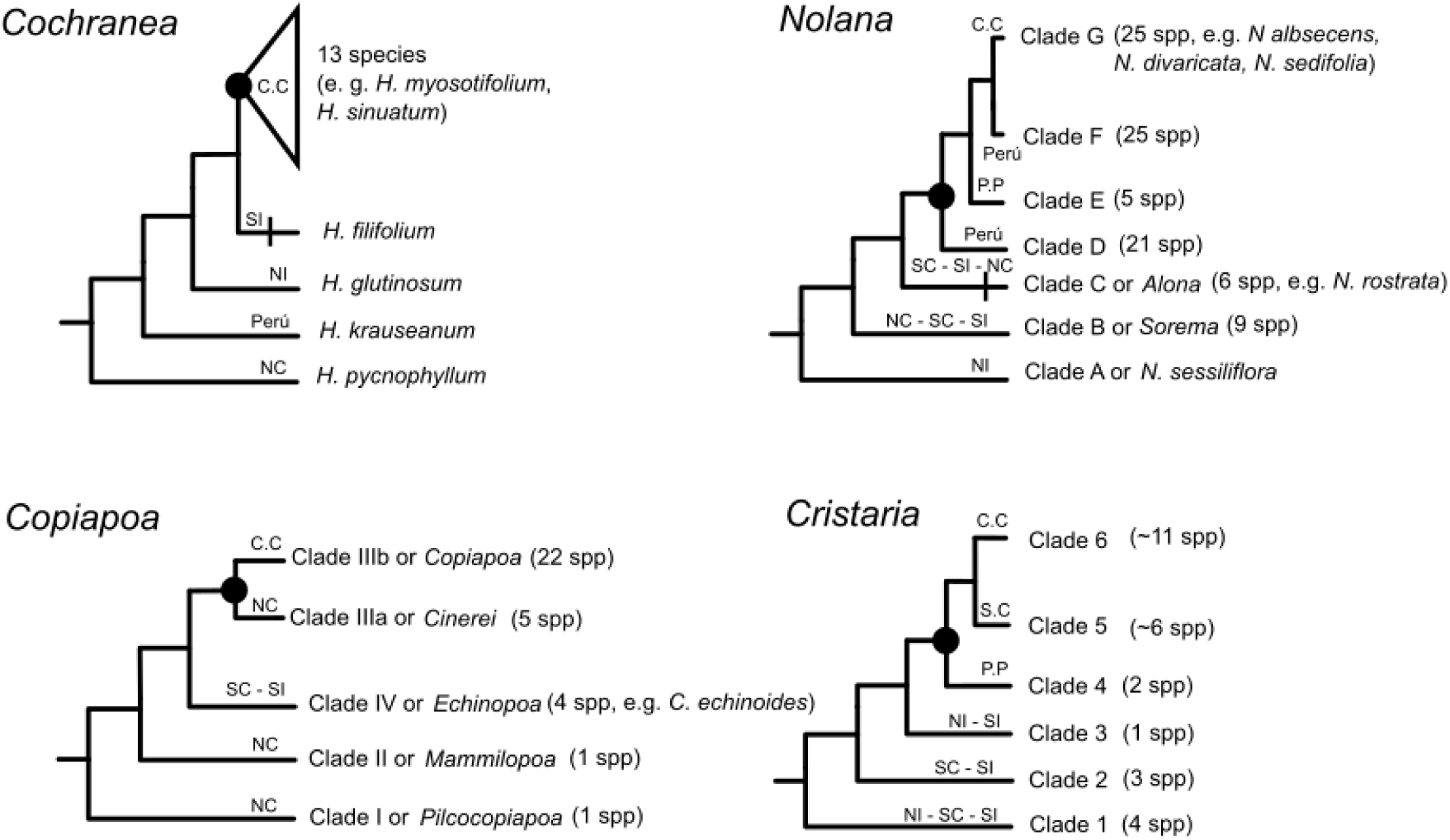
Common evolutionary history of specious plant genera from the Atacama Desert. The simplified topologies of *Cochranea*, *Nolana*, *Copiapoa* and *Cristaria* are shown. Black bars represent the emergence of terete leaves. Black dots represent the ancestor that experienced adaptive radiation during a dry phase. Letters above branches depict species distribution, NI, Northern Interior; NC, Northern Coast; SI, Southern Interior; SC, Southern Coast; P.P, Perú and Prepuna; C.C, Broad Chilean distribution

The high predictive power of the model suggests that, in both scenarios (3 or 4 climatic zones), *Nolana* and *Cochranea* species are generally restricted to a specific climatic zone, mostly either north or south of 27°S. The exceptions included in the analysis were *N. divaricata*, *N. sedifolia*, *N. elegans*, *N. acuminata*, and *H. floridum*. It is highly likely that including more species in the analysis would only reinforce the overall climatic specificity of *Nolana* and *Cochranea* species across the defined zones.

### 3.4 Phylogeny of *Cochranea* and Ancestral Area Reconstruction

The alignment of the five loci had a length of 5.700 base pairs. The topology of the resulting phylogenetic tree is consistent with the literature, corroborating the sequence of basal clades (*H. pycnophyllum* (*H. krauseanum* (*H. glutinosum* (*H. filifolium* (12 species)))) (Fig. S8). The clade that groups *H. filifolium* and the core of species diversity showed lower support than the previous clades, which is probably due to the phylogenetic closeness between *H. filifolium* and *H. glutinosum* (Fig. S8).

The S-DIVA analysis suggests that the ancestral distribution range of *Cochranea* was located north of the 27°S parallel (nodes 31, 30, and 29) and that *H. filifolium* represents a dispersal to the south (Fig. S9). Node 28 suggests that the common ancestor of *H. filifolium* and the core of *Cochranea* diversity was distributed in the Southern Interior (Marginal Desert) and in the Northern Coast. This is consistent in part with biogeographic reconstructions carried out for *Copiapoa* and *Cristaria*, where said common ancestor was distributed south of the Copiapó river (Larridon et al., 2015) and in the Atacama southern pampa (which corresponds mainly to the Southern interior zones in our analysis) (Böhnert et al., 2022).

## 4. Discussion

### 4.1 Community structure of vegetation in a geomorphological transition and habitat differentiation between congeneric species

Desert environments present limitations in water availability, which generates an irregular distribution of plants (Borthagaray et al., 2010; Stanton et al., 2014). Here, plant assemblages were characterized in a site located in the transition between two geomorphological units composing the landscape of the III Region, called “Coastal Cordillera” (high ruggedness) and “Transitional Pampa” (plain) (Novoa et al., 2008). This transitional characteristic gives the study site a greater heterogeneity of water conditions than areas located within a single geomorphological unit. For example, within the Coastal Cordillera south of the Huasco River, indicator species of group 9 such as *E. acida*, *M. miquelii*, and *C. coquimbana* (see section 4.3) grow homogeneously in the landscape occupying entire basins, while in the study site they mainly inhabit the altitudinal range of the fog (Fig. 3, S4). It is likely that, in the transition from the Coastal Cordillera to the plains of the Transitional Pampa (i.e. the study site), the decreased basins and hills density (i.e., decrease in ruggedness) leads to lower water retention in the soil. Consequently, water availability varies markedly depending on the altitudinal position of the fog, which in turn influences the distribution of vegetation formations.

Some community patterns found here were also observed in Paposo (25°S), where the difference in water availability granted by fog is also highly variable in the altitudinal gradient. In Paposo, a peak of species diversity and growth forms is recorded at the upper and lower edges of the fog range (Rundel & Mahu, 1976). In the study site, the group 8 presents the greatest abundance and species richness (Fig. S5), and it is also distributed at the edge of the fog (Figs. 3, S4). Although the tendency to a northern orientation of this group could be the cause of its distinction in the SIMPROF analysis and of the observed abundance and richness, data obtained in Paposo suggest that, in reality, altitude and proximity to fog are the key factors. This pattern seems related to the presence of an ecotone between an habitat with high water availability and a more arid one. This hypothesis contrasts with what has been proposed for Paposo, where the higher diversity of species and life forms at the margins of the fog is said to be caused by less competition among species with the same growth forms, whereas in areas within the fog range competition is stronger (Rundel & Mahu, 1976).

In general, the plant assemblages defined here can be divided into two groups based on slope percentage: slope formations (groups 3, 6, 7, 8, and 9) and dry washes or plain formations (groups 4, 5, and 10) (Figs. 3, S3). In group 4, species richness and abundance comparable to group 9 are observed, and it is possible that these two groups represent formations with the highest water availability, although in the first the water source arrives by runoff and in the second by direct contact with the fog. Although an ecological differentiation between stream species versus slope species is expected, particularly in their root system, (Rundel et al., 1980), several species that are not indicative of any group inhabit both types of formations, such as *Adesmia argent*ea, *Balsamocarpon brevifolium*, and *Cordia decandra* (Table S1). These species usually show clear spatial differentiation when dominant in certain localities, where *A. argentea* and *B. brevifolium* occupy south-facing slopes and *C. decandra* occupies north-facing slopes. It is possible that dry washes are favorable environments for the joint establishment of species with different habitat preferences in terms of slope orientation or fog-derived water requirements. It has been pointed out that congregations of several *Nolana* species may be due to surface runoff transport of fruits coming from different slopes (Dillon, 2023). Consistent with this, the only plot where the three *Cochranea* species were recorded corresponds to group 4. Dry washes have been catalogued as biological and gene flow corridors in desert environments (Gaddis et al., 2016).

The community structure of the study site suggest that on the slopes water availability decreases as a function of distance from the altitudinal range of the fog. Groups 5, 6, and 7 show an increase in the abundance of *C. sphaerica* with altitude, peaking in group 8 and almost disappearing in group 9 (Table S1). Additionally, it is common to observe that vegetation disappears at the foot of the hills (Fig. S10), suggesting a decrease in total abundance as a function of altitude, although this was not significant (Fig. S6). Probably if only slope vegetation formations were considered this relationship would have been clearer. A more exhaustive analysis focused on each species and its altitude-abundance relationship remains pending.

An unexpected finding, not evident during data collection except in the case of *H. myosotifolium* and *H. filifolium*, is the niche segregation pattern among congeneric species observed in the study site, both in *Nolana* and *Cochranea* (Fig. 2, S6). Furthermore, the species *B. brevifolium* and *E. angulatus*, which used to be circumscribed to *Caesalpinia*, also segregate in the multivariate analysis. The results of this study demonstrate that, although some congeneric species can share the same habitat and their ecological differences are not evident at first glance, a sufficiently broad sampling can reveal them. Despite ecological differentiation between *H. myosotifolium* and *H. filifolium*, primarily related to altitude (Fig. S6), both species retain the characteristic of distributing mainly on slopes, and the segregation they exhibit just a few meters apart and on two slopes at the same altitude remains curious (Fig. S2).

### 4.2 Evolutionary history of species-rich plant genera of the Atacama Desert

In the study site, *N. rostrata*, *H. filifolium*, and *C. echinoides* are indicator species of group 9 and inhabit the altitudinal range of the fog, thus preferring habitats with greater water availability (Figs. 2, 3). The fact that they belong to the same basal clade within their respective genera supports the hypothesis of a shared evolutionary history, which is likely even closer between *N. rostrata* and *H. filifolium* given their convergence in leaf morphology (Fig. 6).

**Figure 6.**
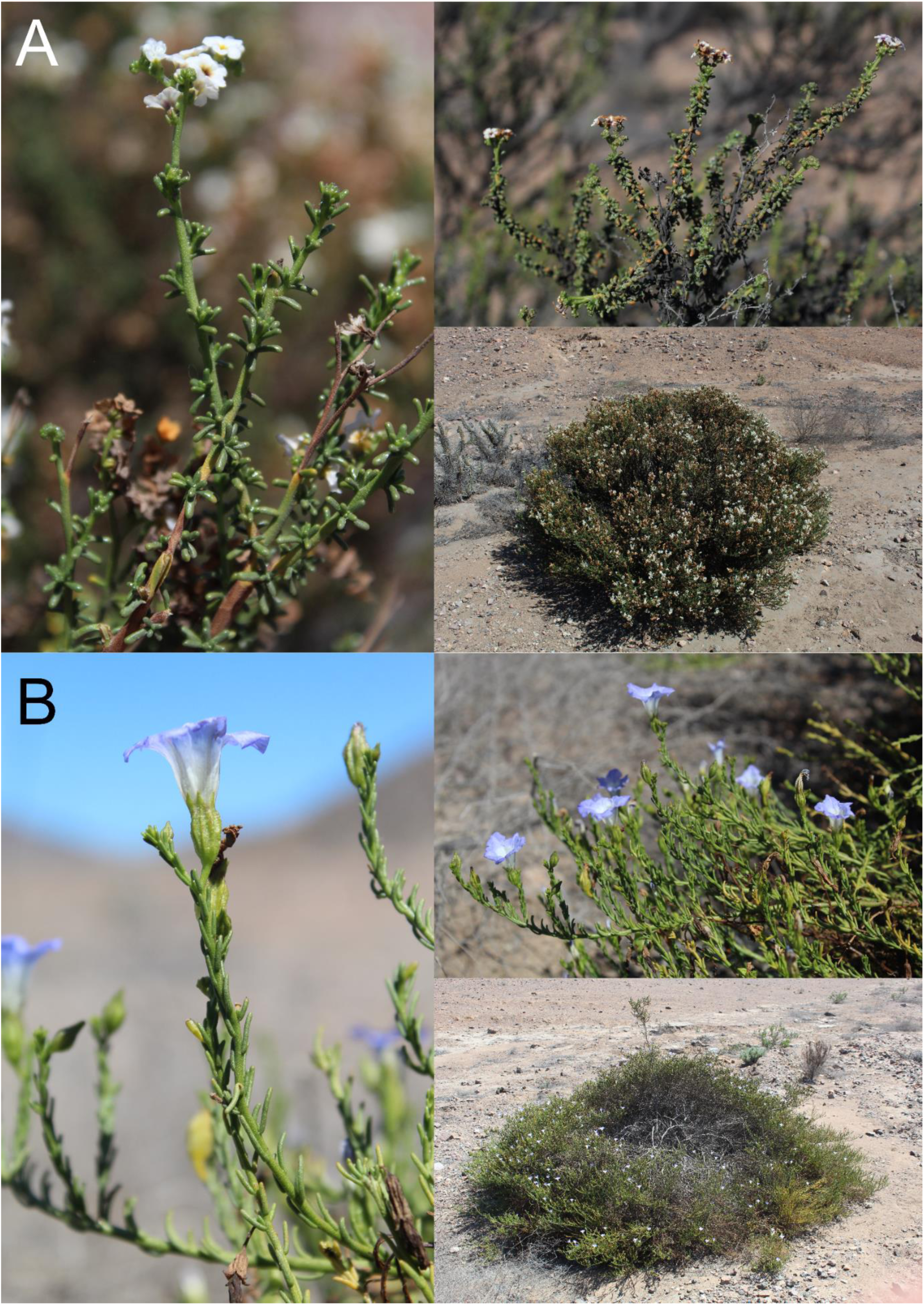
Morphological convergence in *Nolana* and *Cochranea* into terete leaves. A, *Heliotropium filifolium*. B, *Nolana rostrata*.

In *Nolana*, *Cochranea*, *Copiapoa* and *Cristaria*, a similar sequence of phylogenetic divergences is observed, in which analogous branches are composed of species distributed in similar biogeographic areas (Fig. 5). Basal clades are mainly distributed north of parallel 27°S, and more clearly, the clade that is sister to the diversity core in each genus is distributed in the Southern Interior climatic zone in the marginal desert, south of parallel 27°S. Recently diverged species, composing the diversity core of each genera, are broadly distributed in Chile and also in Peru in the case of *Nolana* and *Cristaria*. This phylogenetic pattern reflects a northern migration that started in the marginal desert, and the posterior diversification.

The paleoclimatic history of the Atacama Desert is far from reaching a consensus (Jordan et al., 2014). However, if the evolution of the flora is considered as evidence, it is possible to hypothesize key events, without intending to be precise in terms of their dates. The formation of the hyperarid core led to extinction and vicariance of plant lineages. Its formation may have been caused either by the uplift of the Andes and the consequent increase in the rain shadow effect, or by the abrupt uplift of the continental shelf and the formation of the coastal cliff in northern Chile, an event that occurred between the Late Miocene and early Pliocene (Paskoff, 1979), and which would have limited the entry of moisture from the Pacific Ocean into the interior. This event gains relevance given that the importance of the rain shadow effect in the formation of the hyperarid core has been questioned (Garreaud et al., 2010). Other factors that may also have influenced the formation of the hyperarid core include the cooling of ocean temperatures and the consequent intensification of the thermal inversion due to subsidence (Garreaud et al., 2010).

Probably the vegetation that inhabited the area of the hyperarid core began to become extinct when extreme aridity conditions started, leaving remnants in the surrounding areas. In line with this, *N. sessiliflora* and *H. glutinosum* are basal clades that adapted to survive these extreme conditions, and it is likely that several lineages that today inhabit fog zones also did so in the hyperarid core area prior to the establishment of hyperarid conditions. In *Cochranea*, this vicariant divergence is clear, since the basal species have allopatric distributions (*H. pycnophyllum*: Northern Coast; *H. krauseanum*: Peru; *H. glutinosum*: Northern Interior; *H. filifolium*: Southern Interior) (Fig. 5). The age of this divergence was estimated at around 7.4 million years ago, during the late Miocene (Luebert & Wen, 2008). Later, between 4.6 – 3.79 million years ago during the Pliocene, the adaptive radiation characteristic of the diverse plant genera of the Atacama Desert began (Luebert & Wen, 2008; Dillon et al., 2009; Luebert et al., 2011; Guerrero et al., 2013; Böhnert et al., 2019, 2022). The dominant hypothesis proposes that alternations between short wet and long dry phases during the Plio-Pleistocene (Jordan et al., 2014) led to expansions and contractions in the distribution of plant species, generating speciation through geographic isolation. Another diversification model compatible with these climatic fluctuations is proposed here: the ancestor that underwent adaptive radiation expanded its distribution during a wet phase, from the Marginal Desert to the north, reaching as far as Peru in the case of *Nolana* and *Cristaria* (Fig. 5). At the onset of the following arid phase, there was an increase in the heterogeneity of water availability at a small spatial scale, similar to what is observed in Paposo or in the study site, thus producing adaptative radiation. The resulting phylogenetic pattern results in that the clade from the Marginal Desert (from where the migration began) is basal to the clade that migrated furthest (Peru in *Nolana* and *Cristaria*), and the subsequent derived clades are geographically distributed between the first two, along the Chilean coast. Therefore, probably the migration route was by the coast, and not through the Andes. *Nolana* underwent a second colonization and diversification in Peru (Dillon et al., 2009).

About the speciation events, individuals of this widely distributed common ancestor were exposed to different hydric conditions, resulting in local adaptation events, that is, ecological speciation, where genetic isolation between sets of individuals subjected to different abiotic conditions occurs alongside phenotypic and ecological changes just as happens with speciation mediated by edaphic factors (Nosil et al., 2009; Sakaguchi et al., 2019). Speciation events such as the one just described may have occurred in each humid-arid cycle and are in line with practical and theoretical evidence that questions whether population isolation and associated genetic differentiation alone are sufficient to produce speciation (Maturana-Romesin & Mpodozis, 2000; Sakaguchi et al., 2019; Singhal et al., 2022; Castro et al., 2023).

The diversification model proposed here, in which a common ancestor expands its distribution and subsequently undergoes cladogenesis due to increased hydric heterogeneity at the landscape scale, implies that phylogenetically related species would tend to inhabit the same climatic zone. This appears to be the case in the Atacama Desert: a north–south division is observed, where species north of the Copiapó River are generally more closely related to each other than to those in the south, and vice versa. This occurs with the species of *Alona* (Tu et al., 2008; Dillon et al., 2009), and may also occur in Clade G of *Nolana*, although phylogenetic relationships still need to be resolved. In *Cochranea*, there are two diversity peaks located north and south of the 27°S parallel (Luebert & Wen, 2008), but again, phylogenetic relationships must be clarified. This north–south division is also observed in *Eulychnia* (Larridon et al., 2018; Merklinger et al., 2021) and in some clades of *Cristaria* (Clade 6a, 6b).

The Copiapó River Valley is an important phylogeographic barrier (Ossa et al., 2013), as it represents a flat area that divides the Coastal Cordillera in two. It also marks a unique geomorphological transition in Chile: north of the 27°S parallel lie the Altiplanic landscape and the “Pampa ondulada” where the hyperarid core is located (Novoa et al., 2008). The 27°S parallel also marks the boundary between arid and hyperarid climates (Luebert & Pliscoff, 2006). Therefore, the area of influence of the Copiapó River constitutes a climatic and geographic barrier that may have favored the emergence of parallelisms (Larridon et al., 2015), reinforcing the status of the Atacama Desert as a natural laboratory.

Within the diversity of leaf forms present in *Nolana* and in *Cochranea* (Luebert, 2013; Dillon, 2023), the convergence toward a terete morphology in *H. filifolium* and *N. rostrata* (and in *Alona* in general) has gone unnoticed until now. These species differ in that in *H. filifolium*, the leaves are grouped in fascicles, whereas in *N. rostrata*, they grow solitarily. In both species, the leaves are narrow and abundant, and the branches are positioned very close to one another (Fig. 6), which suggests that this morphology has evolved to optimize the capture of fog microdroplets, allowing their subsequent runoff toward the root system (Martorell & Ezcurra, 2007; Vogel & Müller-Doblies, 2011). This latter point is consistent with the root arrangement reported in some *Cochranea* species, where the roots are superficial and spread beneath the shrub canopy (Rundel et al., 1980; Olivares & Squeo, 1999). Other morphological convergences in the Atacama flora include *Malesherbia fasciculata* (Malesherbiaceae), which adopts the growth habit of species in the genus *Polyachyrus* (Asteraceae) (Bull-Hereñu, 2020), and the foliar similarity between *H. sinuatum* and *Pleocarphus revolutus* (Asteraceae).

## 5. Conclusion

The six plant assemblages present in the study site are differentially distributed along the altitudinal gradient. The congeneric species of *Nolana* and *Cochranea* showed a pattern of niche segregation, which supports the hypothesis of ecological speciation events as being responsible for the diversity of the flora in the Atacama Desert – Peru. The evolutionary history of some diverse genera converges in terms of timing and biogeography, and reflects a phylogenetic pattern in which the clade from the Marginal Desert (from where the migration began) is basal to the one that migrated furthest (Peru in *Nolana* and *Cristaria*), and the subsequent derived clades are geographically distributed between the first two. This phylogenetic pattern likely reflects that, after the northward migration, aridity progressively advanced southward, enhancing the evolution and divergence of the northernmost populations first. A good question refers to the extent of the presence of the ancestor that migrated north and diversified, that is, whether it was only along the coast or also included inland areas. It is also necessary to explain why the ancestor from the Marginal Desert migrate and diversified while the other basal clades did not, something that may be related to his fog-adapted ecology, evidenced by the leaf morphology of *Alona* and *H. filifolium*. The ecological and evolutionary similarities between *N. rostrata* and *H. filifolium* are a clue that helped to discover the common evolutionary history of the Atacama flora and present an opportunity to understand through which means the leaf convergence occurred. The Atacama Desert constitutes a natural laboratory, in which its flora has diversified within the same evolutionary context, and therefore will allow valuable comparisons to understand evolutionary phenomena. For example, to answer why some genera are more diverse than others.

When the fog vegetation of northern Chile is mentioned, people mistakenly think about the relict hygrophilous forest of Fray Jorge (30°39’S) or the bloom of annual herbs called “Desierto Florido” that occur during years with El Niño precipitations. It is evident that there is a general lack of knowledge about this biodiversity and its threats, which lie in large-scale mining, goat and mule livestock, and human housing expansion. The conservation status of *H. filifolium* is Vulnerable due to its small area of occupancy (692 km²) with only 9 known localities and therefore in the future we could lose the opportunity to continue learning from this study model.

## Acknowledgments

This study is largely based on the work of Federico Luebert and Michael O. Dillon, to whom I am grateful. Thanks to Laurent Hardion for providing valuable comments on the manuscript. Special thanks to those who made this research possible: Sergio Durán, Carlos Miranda, Stefany Martínez, Felipe Roa, Rodrigo Petit, Pilar Mena, Victoria Téllez, and Francisca Reveco, for their collaboration in data collection, species identification, and for their advice.

