## Supplementary material 1 for "Exploring the fog vegetation in the marginal Atacama Desert: common evolutionary history of species-rich genera and morphological convergence in *Nolana* and *Heliotropium* sect. *Cochranea*"

**Appendices**

Table S1. Floristic diversity at the study site

| **Species** | **Family** | **Tribe** | **Origin** | **Conservation status** | **Mean abundance per Simprof group** | | | | | | | | **Total** |
| --- | --- | --- | --- | --- | --- | --- | --- | --- | --- | --- | --- | --- | --- |
|  |  |  |  |  | **3 (n=17)** | **4 (n=49)** | **5 (n=6)** | **6 (n=5)** | **7 (n=8)** | **8 (n=8)** | **9 (n=37)** | **10 (n=12)** |  |
| *Encelia canescens* Lam. var. *canescens* | Asteraceae | Heliantheae | Native | - | 4,94 | 37,37 | 1,67 | 11,4 | 1,38 | 8 | 2,05 | 0,5 | 2139 |
| *Cumulopuntia sphaerica* (C.F. Först.) E.F. Anderson | Cactaceae | Tephrocataceae | Native | Least Concern (LC) | 2,82 | 3,95 | 5,33 | 12,8 | 20,38 | 46 | 1,51 | 0,25 | 928 |
| *Eulychnia acida* Phil. | Cactaceae | Notocactaceae | Endemic | Least Concern (LC) | 1,76 | 1,2 | 0,5 | 4,6 | 1,63 | 4,63 | 11,68 | 0,67 | 605 |
| *Miqueliopuntia miquelii* (Monv.) F. Ritter | Cactaceae | Opuntieae | Endemic | Least Concern (LC) | 1,06 | 1,61 | 0,83 | 0,8 | 3,5 | 1,88 | 10,97 | 0,17 | 557 |
| *Heliotropium filifolium* (Miers) I.M. Johnst. | Boraginaceae | Boragineae | Endemic | Vulnerable (VU) | 0 | 1,84 | 0,67 | 0,2 | 1,25 | 2,25 | 10,97 | 0,25 | 532 |
| *Erythrostemon angulatus* (Hook. & Arn.) Gagnon & G.P Lewis | Fabaceae | Caesalpinieae | Endemic | - | 2,35 | 4,47 | 0,17 | 0,6 | 0,25 | 8,67 | 1,46 | 0,33 | 392 |
| *Heliotropium myosotifolium* (A. DC.) Reiche | Boraginaceae | Boragineae | Endemic | - | 12,53 | 0,37 | 0,5 | 1,2 | 0,87 | 0,5 | 0,03 | 0 | 252 |
| *Adesmia argentea* Meyen | Fabaceae | Dalbergieae | Endemic | - | 3,82 | 2,37 | 0,33 | 4.4 | 0,75 | 2,75 | 0,14 | 0 | 238 |
| *Balbisia peduncularis* (Lindl.) D. Don | Francoaceae | - | Native | - | 1,35 | 0.9 | 0 | 0 | 0,13 | 2 | 2,03 | 0 | 159 |
| *Fagonia chilensis* Hook. & Arn. | Zygophyllacae | Heliantheae | Native | - | 1,12 | 0,71 | 0,5 | 0,6 | 0 | 4,63 | 0,32 | 1,8 | 131 |
| *Nolana albescens* (Phil.) I.M. Johnst. | Solanaceae | Nolaneae | Endemic | - | 1,88 | 0,65 | 0 | 0 | 0 | 2,88 | 1,18 | 0 | 131 |
| *Krameria cistoidea* Hook. & Arn. | Krameriacae | - | Endemic | Least Concern (LC) | 0,34 | 0,76 | 0 | 0,8 | 0 | 2 | 0.81 | 0 | 91 |
| *Tetragonia angustifolia* Barnéoud | Aizoaceae | - | Endemic | - | 0,17 | 0,06 | 0 | 0 | 0,38 | 2 | 1,68 | 0 | 87 |
| *Frankenia chilensis* K. Presl | Frankeniaceae |  | Native | - | 0,24 | 0,94 | 0 | 0,4 | 0,13 | 0,25 | 0,61 | 0,33 | 82 |
| *Tillandsia landbeckii* Phil. | Bromeliaceae | - | Native | - | 0 | 0 | 0 | 0 | 0 | 0 | 2,08 | 0 | 77 |
| *Nolana divaricata* (Lindl.) I.M. Johnst. | Solanaceae | Nolaneae | Endemic | - | 0,17 | 1,24 | 0,33 | 0 | 0 | 0,5 | 0 | 0,33 | 74 |
| *Balsamocarpon brevifolium* Clos | Fabaceae | Caesalpinieae | Endemic | Endangered (EN) | 0,59 | 0,1 | 0,17 | 1 | 0,38 | 2,25 | 1,03 | 0 | 71 |
| *Nolana rostrata* (Lindl.) Miers ex Dunal | Solanaceae | Nolaneae | Endemic | - | 0,06 | 0,08 | 0 | 0 | 0,13 | 0 | 1,7 | 0 | 69 |
| *Nolana sedifolia* Poepp. | Solanaceae | Nolaneae | Endemic | - | 0,35 | 0,38 | 0 | 0,2 | 0,13 | 1,88 | 0,68 | 0,08 | 68 |
| *Senecio myriophyllus* Phil. | Asteraceae | Senecioneae | Endemic | - | 0,24 | 0,65 | 0 | 0 | 0,16 | 0,24 | 0,57 | 0 | 60 |
| *Atriplex deserticola* Phil. | Amaranthaceae | Atripliceae | Native | - | 0 | 0,28 | 0 | 0 | 0 | 0,75 | 0,95 | 0 | 55 |
| *Cordia decandra* Hook. & Arn. | Boraginaceae |  | Endemic | Near Threatened (NT) | 0,12 | 0,67 | 0,17 | 0 | 0 | 0,75 | 0,35 | 0 | 55 |
| *Lycium bridgesii* (Miers) R.A. Levin, Jill S. Mill. & G. Bernardello | Solanaceae | Lycieae | Endemic | - | 0,35 | 0,47 | 0 | 0,6 | 0,25 | 0,5 | 0,35 | 0 | 51 |
| *Pleurophora pungens* D. Don. | Lythraceae | - | Endemic | - | 0,17 | 0,65 | 0 | 0,4 | 0 | 0,5 | 0 | 0 | 41 |
| *Senna cumingii* (Hook. & Arn.) H.S. Irwin & Barneby var. *cumingii* | Fabaceae | Cassieae | Endemic | - | 0,06 | 0,65 | 0 | 0 | 0,13 | 0,25 | 0,57 | 0 | 39 |
| *Copiapoa echinoides* (Lem. Ex Salm-Dyck) Britton & Rose | Cactaceae | Notocactaceae | Endemic | Vulnerable (VU) | 0,05 | 0,02 | 0 | 0 | 0,25 | 0,38 | 0,76 | 0 | 35 |
| *Galenia pubescens* (Eckl. & Zeyh.) Druce | Aizoaceae |  | Introduced | - | 0 | 0,67 | 0 | 0 | 0 | 0 | 0 | 0 | 33 |
| *Skytanthus acutus* Meyen. | Apocynaceae | Plumerieae | Endemic | - | 0 | 0,2 | 0 | 0 | 0 | 0 | 0 | 1,83 | 32 |
| *Heliotropium sinuatum* (Miers) I.M. Johnst. | Boraginaceae | Boragineae | Endemic | - | 0 | 0,4 | 0 | 0 | 0 | 0,14 | 0,08 | 0 | 26 |
| *Errazurizia multifoliolata* (Clos) I.M. Johnst. | Fabaceae | Amorpheae | Endemic | - | 0,59 | 0 | 0 | 0 | 0 | 0.86 | 0.03 | 0 | 18 |
| *Oxalis gigantea* Barnéoud | Oxalidaceae | - | Endemic | - | 0 | 0,34 | 0 | 0 | 0 | 0 | 0,03 | 0 | 18 |
| *Cheilanthes mollis* (Kunze) C. Presl | Pteridaceae | Trichomaneae | Native | Least Concern (LC) | 0 | 0 | 0 | 0,2 | 0 | 1,25 | 0,16 | 0 | 17 |
| *Nicotiana glauca* Graham | Solanaceae | Nicotianeae | Native | - | 0,56 | 0,1 | 0 | 0 | 0 | 0 | 0 | 0 | 15 |
| *Argylia radiata* (L.) D. Don | Bignoniaceae | Tecomeae | Native | - | 0 | 2,45 | 0 | 0 | 0 | 0 | 0 | 0 | 12 |
| *Ophryosporus triangularis* Meyen | Asteraceae | Eupatorieae | Endemic | - | 0 | 0,02 | 0 | 0,2 | 0 | 0,5 | 0,1 | 0 | 10 |
| *Polyachyrus poeppigii* Kuntze ex Less. subsp. *poeppigii* | Asteraceae | Nassauvieae | Native | - | 0 | 0 | 0 | 0 | 0 | 0 | 0,24 | 0 | 9 |
| *Centaurea chilensis* Hook. & Arn. var. *chilensis* | Asteraceae | Cardueae | Endemic | - | 0 | 0 | 0 | 0 | 0,16 | 0 | 0,19 | 0 | 8 |
| *Spinoliva ilicifolia* (Hook. & Arn.) G.Sancho subsp*. ilicifolia* | Asteraceae | Nassauvieae | Endemic | - | 0 | 0,08 | 0 | 0 | 0 | 0 | 0,1 | 0 | 8 |
| *Ephedra gracilis* Phil. Ex Stapf | Ephedraceae |  | Endemic | - | 0 | 0,81 | 0 | 0 | 0 | 0 | 0,08 | 0 | 7 |
| *Lycium minutifolium* J. Remy | Solanaceae | Lycieae | Endemic | - | 0,06 | 0,06 | 0 | 0,2 | 0 | 0 | 0 | 0 | 5 |
| *Solanum remyanum* Phil. | Solanaceae | Solaneae | Endemic | - | 0,06 | 0,02 | 0 | 0,4 | 0 | 0,13 | 0,16 | 0 | 5 |
| *Adesmia eremophila* Phil. | Fabaceae | Dalbergieae | Endemic | - | 0,05 | 0,02 | 0 | 0,02 | 0 | 0 | 0 | 0 | 3 |
| *Flourensia thurifera* (Molina) DC. | Asteraceae | Heliantheae | Endemic | - | 0 | 0 | 0 | 0 | 0,13 | 0 | 0,03 | 0 | 2 |
| *Tropaeolum tricolor* Sweet | Tropaolaceae | - | Endemic | - | 0 | 0 | 0 | 0 | 0 | 0 | 0,03 | 0,08 | 2 |
| *Tweedia birostrata* (Hook. & Arn.) Hook. & Arn. | Apocynaceae | Asclepiadeae | Endemic | - | 0,06 | 0,02 | 0 | 0,2 | 0 | 0 | 0 | 0 | 2 |
| *Adiantum chilense* Kaulf. var. chilense | Pteridaceae | Adianteae | Native | Least Concern (LC) | 0 | 0 | 0 | 0 | 0 | 0 | 0.02 | 0 | 1 |
| *Alstroemeria* sp. | Alstroemeriaceae | Alstroemerieae | - | - | - | - | - | - | - | - | - | - | - |
| *Atriplex semibaccata* R. Br. | Amaranthaceae | Atripliceae | Introduced | - | - | - | - | - | - | - | - | - | - |
| *Chaetanthera glabrata* (DC.) F. Meigen | Asteraceae | Mutisieae | Endemic | - | - | - | - | - | - | - | - | - | - |
| *Chenopodium* sp. | Amaranthaceae | - | - | - | - | - | - | - | - | - | - | - | - |
| *Chorizanthe commissuralis* J. Remy | Polygonaceae | - | Native | - | - | - | - | - | - | - | - | - | - |
| *Cistanthe longiscapa* (Barnéoud) Carolin ex Hershkovitz | Montiaceae | - | Endemic | - | - | - | - | - | - | - | - | - | - |
| *Convolvulus sp.* | Convolvulaceae | Convolvuleae | - | - | - | - | - | - | - | - | - | - | - |
| *Cristaria glaucophylla* Cav. var*.* *glaucophylla* | Malvaceae | - | Endemic | - | - | - | - | - | - | - | - | - | - |
| *Cristaria gracilis* Gay | Malvaceae | - | Native | - | - | - | - | - | - | - | - | - | - |
| *Cruckshanksia pumila* Clos | Rubiaceae | Coussareeae | Endemic | - | - | - | - | - | - | - | - | - | - |
| *Cryptantha* sp. | Boraginaceae | Eritrichieae | - | - | - | - | - | - | - | - | - | - | - |
| *Cuscuta chilensis* Ker Gawl. | Convolvulaceae | Cuscuteae | Native | - | - | - | - | - | - | - | - | - | - |
| *Dioscorea* sp. | Dioscoreaceae | - | - | - | - | - | - | - | - | - | - | - | - |
| *Erodium cicutarium* (L.) L’Hér. Ex Aiton | Geraniaceae | - | Introduced |  | - | - | - | - | - | - | - | - | - |
| *Helenium atacamense* Cabrera | Asteraceae | Helenieae | Endemic | - | - | - | - | - | - | - | - | - | - |
| *Hoffmansseggia* sp. | Fabaceae | Caesalpinieae | - | - | - | - | - | - | - | - | - | - | - |
| *Homalocarpus* sp. | Apiaceae | - | - | - | - | - | - | - | - | - | - | - | - |
| *Lepidium* sp. | Brassicaceae | - | - | - | - | - | - | - | - | - | - | - | - |
| *Leucocoryne coronata* Ravenna | Amaryllidaceae | Gillesieae | Endemic | - | - | - | - | - | - | - | - | - | - |
| *Loasa* sp. | Loasaceae | - | - | - | - | - | - | - | - | - | - | - | - |
| *Malesherbia humilis* Poepp. var. *humilis* | Malesherbiaceae | - | Native | - | - | - | - | - | - | - | - | - | - |
| *Menonvillea* sp. | Brassicaceae | - | - | - | - | - | - | - | - | - | - | - | - |
| *Mesembryanthemum crystallinum* L. | Aizoaceae | - | Introduced | - | - | - | - | - | - | - | - | - | - |
| *Mirabilis* sp. | Nyctaginaceae | Nyctegineae | - | - | - | - | - | - | - | - | - | - | - |
| *Oziroë biflora* (Ruiz & Pav.) Speta | Asparagaceae | Oziroeeae | Native | - | - | - | - | - | - | - | - | - | - |
| *Pectocarya dimorpha* (I.M. Johnst.) I.M. Johnst. | Boraginaceae | - | Endemic | - | - | - | - | - | - | - | - | - | - |
| *Perityle emoryi* Torr. | Asteraceae | Perityleae | Native | - | - | - | - | - | - | - | - | - | - |
| *Plantago hispidula* Ruiz & Pav. | Plantaginaceae | - | Endemic | - | - | - | - | - | - | - | - | - | - |
| *Schinus areira* L. | Anacardiaceae | - | Native | - | - | - | - | - | - | - | - | - | - |
| *Sisyrinchium azureum* Phil. | Iridaceae | Sisyrinchieae | Native | - | - | - | - | - | - | - | - | - | - |
| *Tetragonia* cf. *pedunculata* Phil. | Aizoaceae | - | Endemic | Vulnerable (VU) | - | - | - | - | - | - | - | - | - |
| *Viola* sp. | Violaceae | Violeae | - | - | - | - | - | - | - | - | - | - | - |

Table S2. Taxa and GenBank accession of sequences used to reconstruct *Cochranea* phylogeny.

| **Taxon** | **Voucher** | ***ITS1, 5.8S, ITS2*** | ***rps16*** | ***ndhF*** | ***trnL-trnF*** | ***trnS-trnG*** |
| --- | --- | --- | --- | --- | --- | --- |
| ***Cordia*** |  |  |  |  |  |  |
| *Cordia decandra* Hook. & Arn. | Luebert et al. 1873 | EF688903.1 | EF689005.1 | EF688954.1 | EF688851.1 | HQ286105.1 |
| ***Heliotropium*** |  |  |  |  |  |  |
| *Heliotropium curassavicum* L. | Luebert et al. 2521 | EF688896.1 | EF688999.1 | EF688949.1 | EF688843.1 | HQ286080.1 |
| *Heliotropium pycnophyllum* Phil. | Luebert et al. 2813 | EF688868.1 | EF688971.1 | EF688920.1 | EF688815.1 | HQ286058.1 |
| *Heliotropium krauseanum* Fedde | Dillon 8779 | EF688894.1 | EF688997.1 | EF688947.1 | EF688841.1 | HQ286052.1 |
| *Heliotropium glutinosum* Phil. | Luebert et al. 1970 | EF688885.1 | EF688988.1 | EF688938.1 | EF688832.1 | HQ286050.1 |
| *Heliotropium filifolium* (Miers) I. M. Johnston | Luebert et al. 1973 | EF688882.1 | EF688985.1 | EF688935.1 | EF688829.1 | HQ286048.1 |
| *Heliotropium inconspicuum* Reiche | Luebert et al. 2801 | EF688891.1 | EF688994.1 | EF688944.1 | EF688838.1 | HQ286051.1 |
| *Heliotropium stenophyllum* Hook. & Arn. | Luebert et al. 1990 | EF688899.1 | EF689001.1 | EF688950.1 | EF688847.1 | HQ286060.1 |
| *Heliotropium philippianum* I. M. Johnston | Luebert et al. 2124 | EF688887.1 | EF688990.1 | EF688940.1 | EF688834.1 | HQ286057.1 |
| *Heliotropium taltalense* (Phil.) I. M. Johnston | Luebert et al. 2083 | EF688889.1 | EF688992.1 | EF688942.1 | EF688836.1 | HQ286061.1 |
| *Heliotropium megalanthum* I. M. Johnston | Luebert et al. 2165 | EF688876.1 | EF688979.1 | EF688929.1 | EF688823.1 | HQ286055.1 |
| *Heliotropium chenopodiaceum* (DC.) Clos | Luebert et al. 2501 | EF688872.1 | EF688975.1 | EF688924.1 | EF688819.1 | HQ286046.1 |
| *Heliotropium myosotifolium* (DC.) Reiche | Luebert et al. 2011 | EF688878.1 | EF688981.1 | EF688931.1 | EF688825.1 | HQ286056.1 |
| *Heliotropium linariifolium* Phil. | Luebert et al. 2054 | EF688892.1 | EF688995.1 | EF688945.1 | EF688839.1 | HQ286053.1 |
| *Heliotropium sinuatum* (Miers) I. M. Johnston | Luebert et al. 1972 | EF688881.1 | EF688984.1 | EF688934.1 | EF688828.1 | HQ286059.1 |
| *Heliotropium longistylum* Phil. | Luebert et al. 1971 | EF688883.1 | EF688986.1 | EF688936.1 | EF688830.1 | HQ286054.1 |
| *Heliotropium floridum* Clos | Luebert et al. 1974 | EF688884.1 | EF688987.1 | EF688937.1 | EF688831.1 | HQ286049.1 |
| *Heliotropium eremogenum* I. M. Johnston | Luebert et al. 2575 | EF688865.1 | EF688968.1 | EF688917.1 | EF688812.1 | HQ286047.1 |

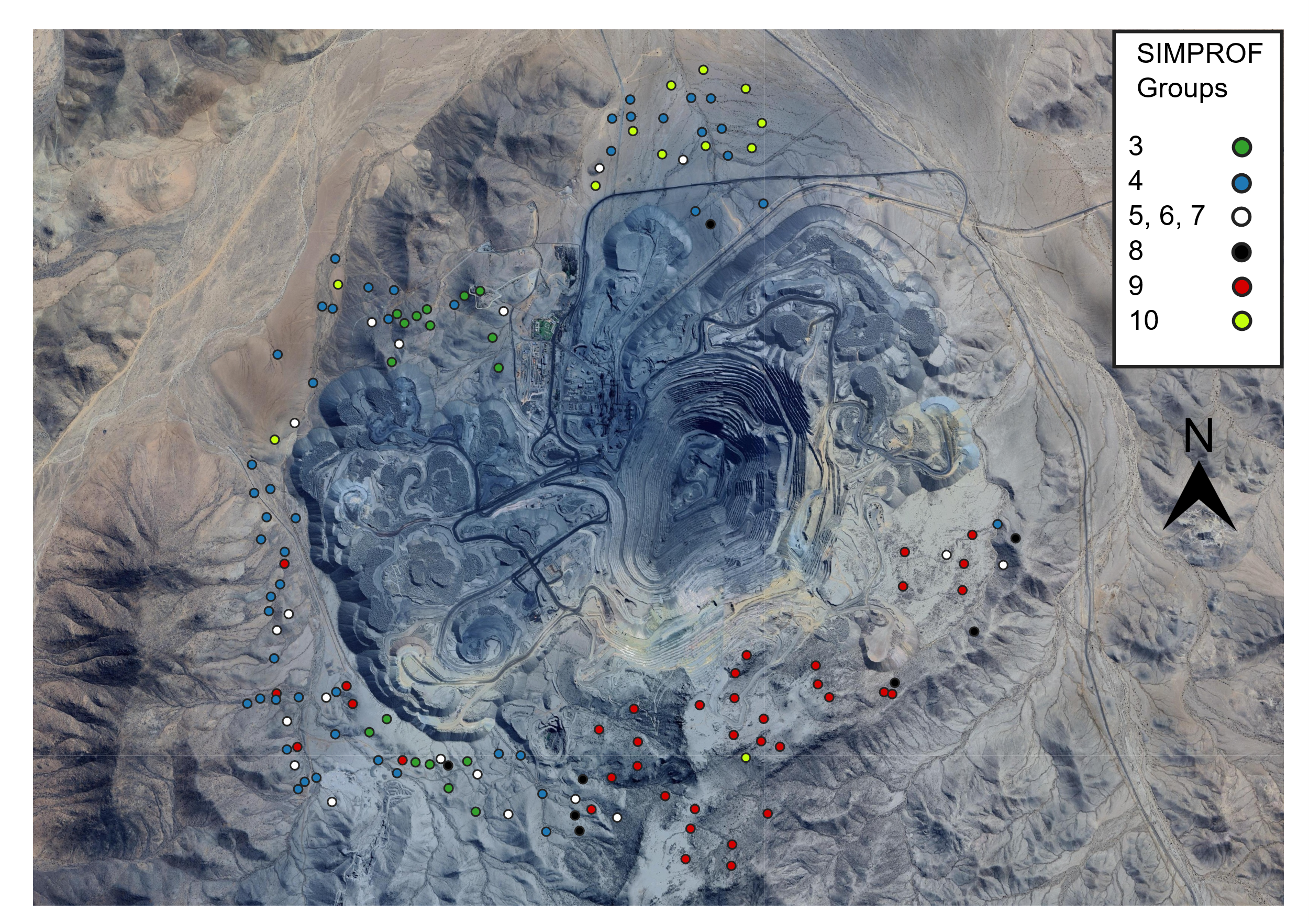

Figure S1. Locations of vegetation plots at the study site and their assignation to Simprof groups.
