## Supplementary material 2 for "Exploring the fog vegetation in the marginal Atacama Desert: common evolutionary history of species-rich genera and morphological convergence in *Nolana* and *Heliotropium* sect. *Cochranea*"

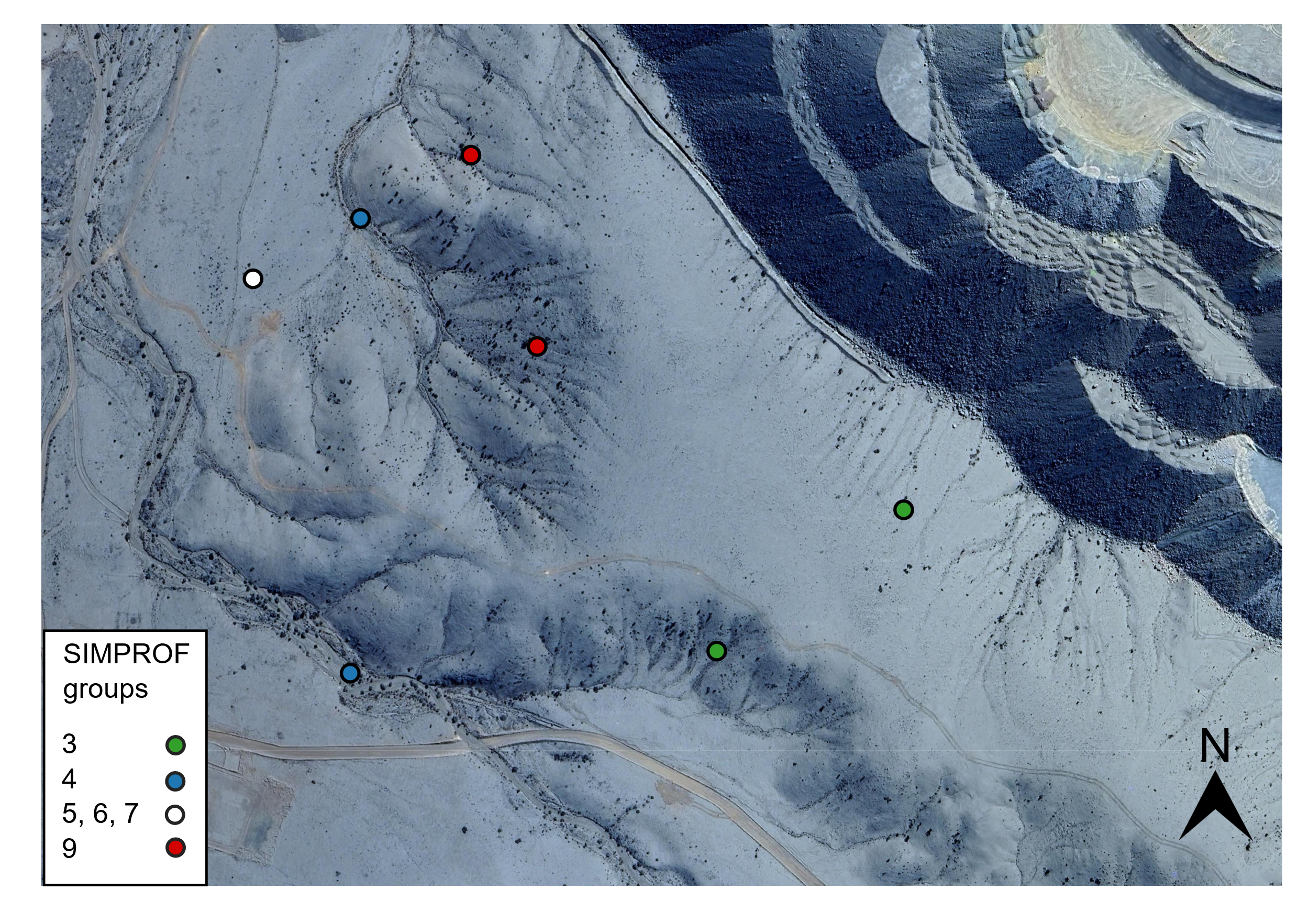

Figure S2. Spatial segregation of *H. filifolium* (Group 9) and *H. myosotifolium* (Group 3).

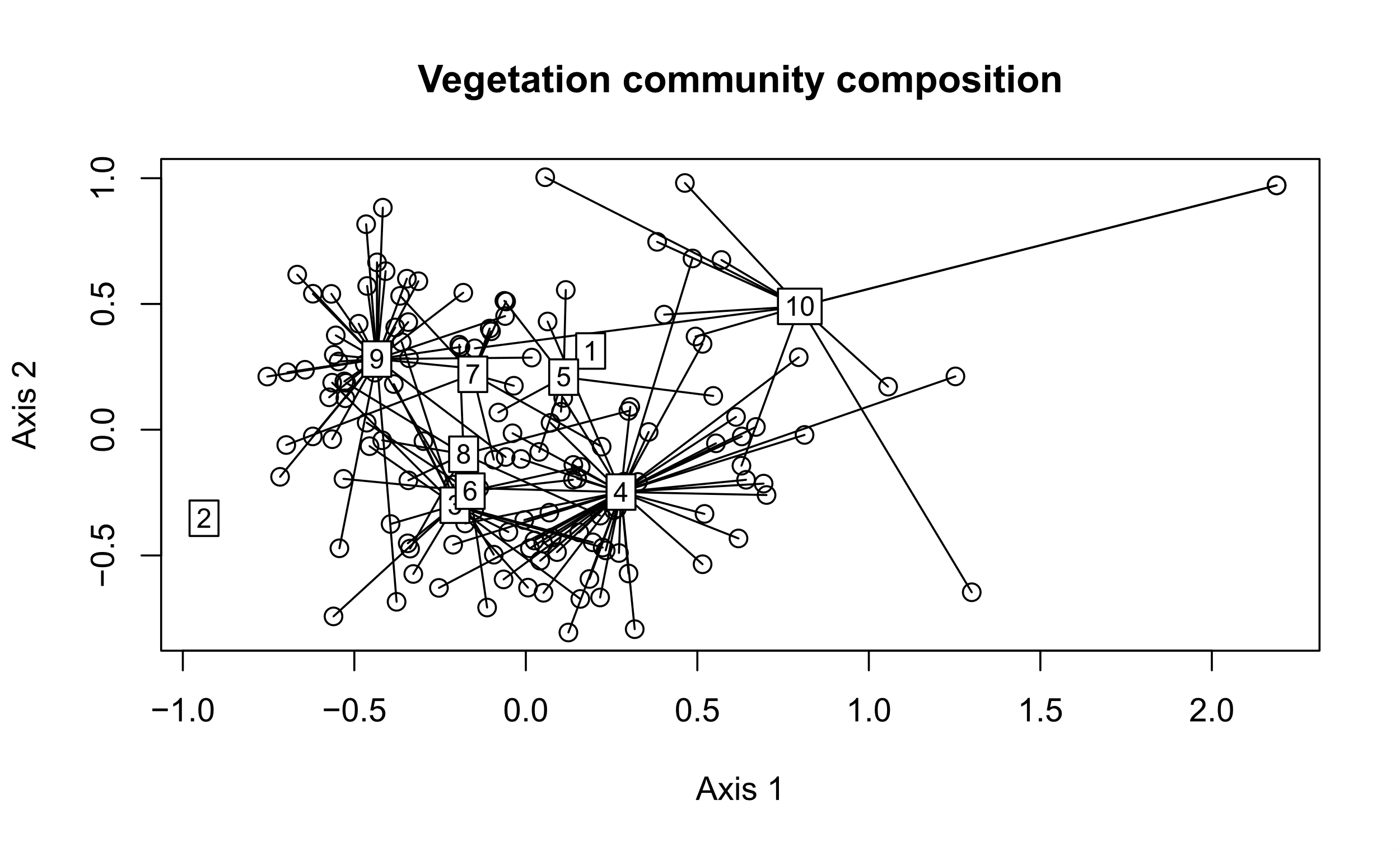

Figure S3. nMDS showing distribution of 144 survey plots based on Bray-curtis dissimilarity. Grouping is based on SIMPROF analysis.

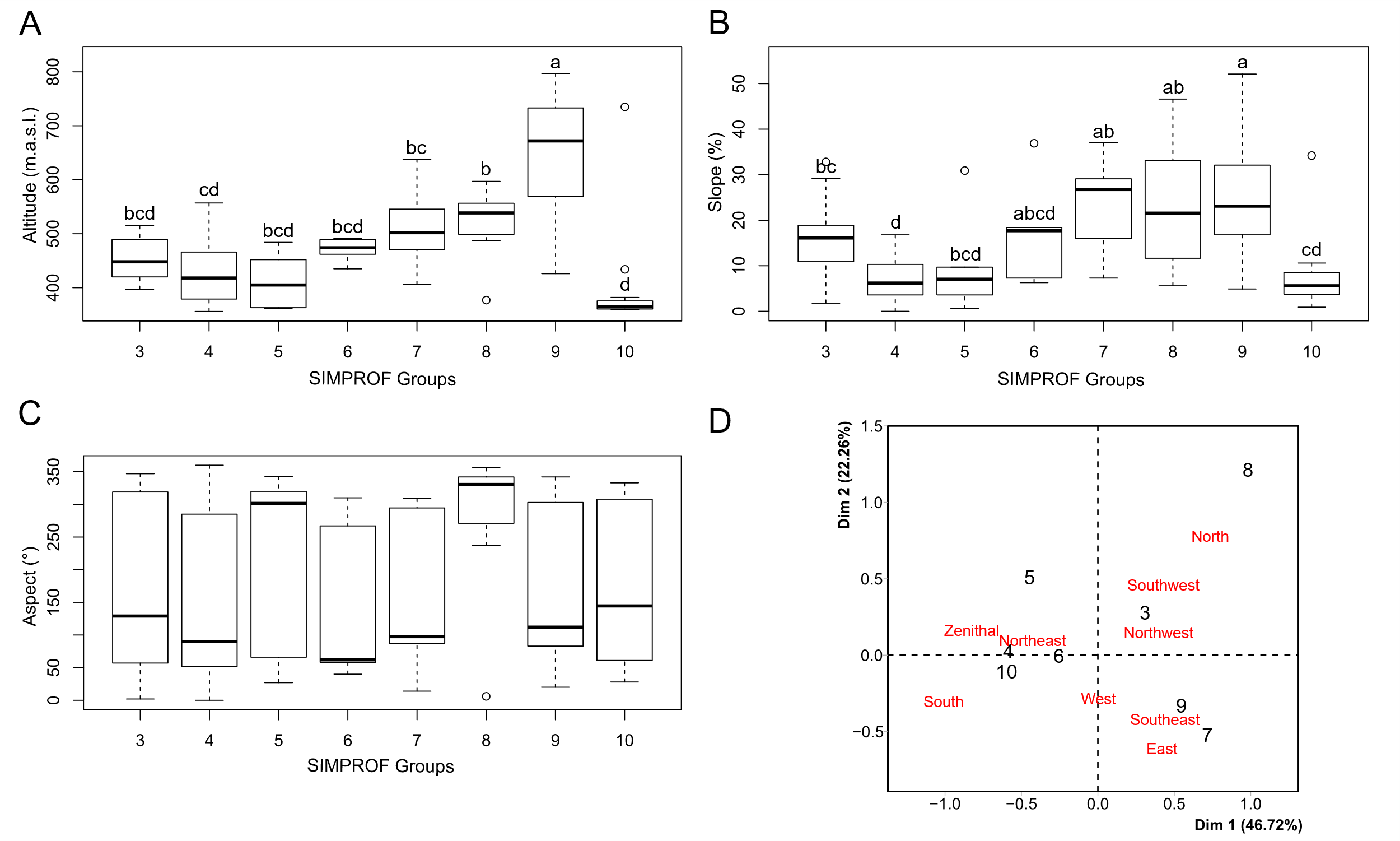

Figure S4. Box-and.whiskers plots for topographic variables of 144 vegetation survey plots. Significant differences (ANOVA and Kruskal Wallis test, Tukey and Wilcoxcon test, respectively) are indicated by different letters above the boxes. A, Altitude. B, Slope. C, Aspect (orientation). D, Correspondence Analysis (CA) of aspect variable transformed to qualitative variable.

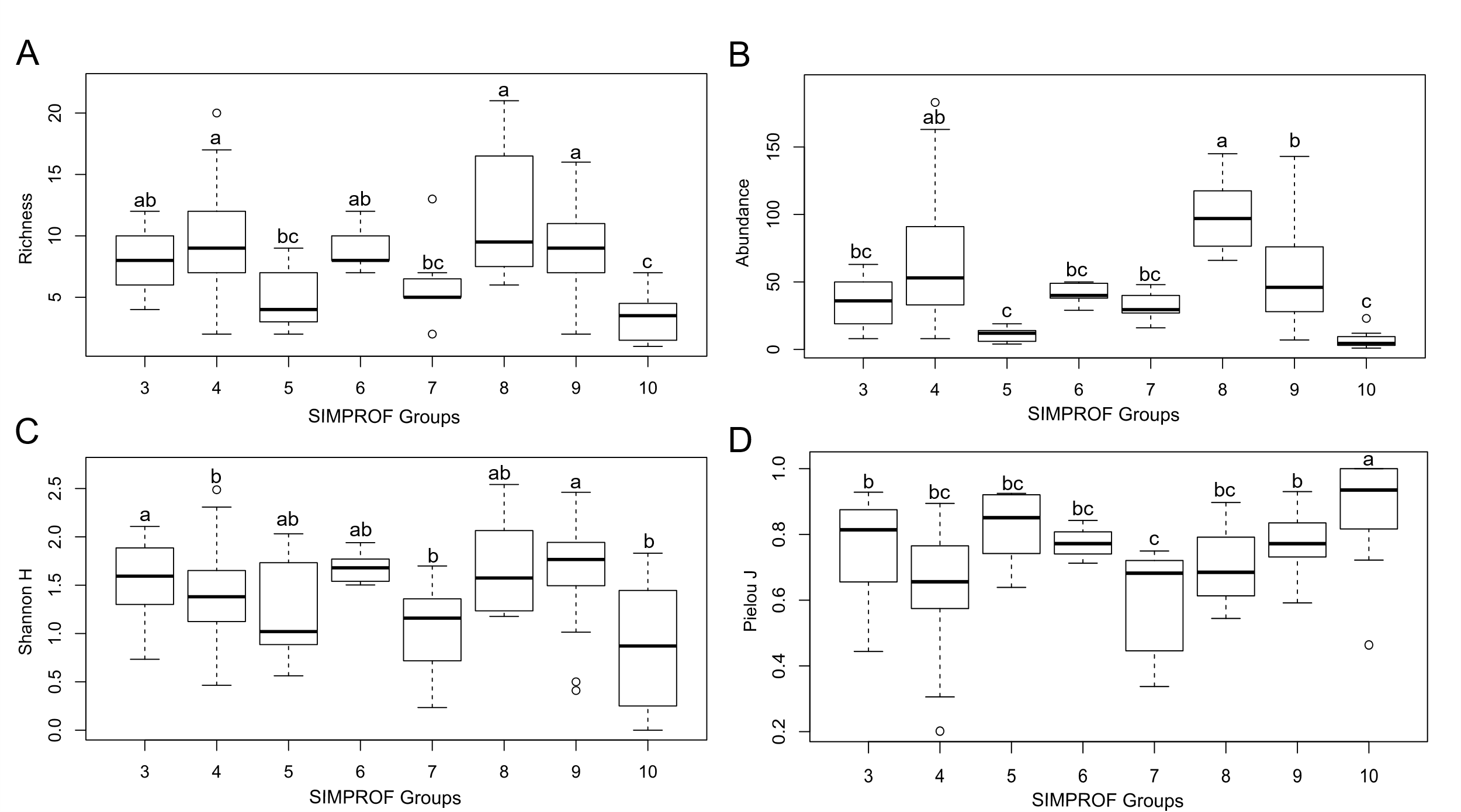

Figure S5. Box-and.whiskers plots for community indices of 144 vegetation survey plots. Significant differences (ANOVA and Kruskal Wallis test, Tukey and Wilcoxcon test, respectively) are indicated by different letters above the boxes. A, Species richness. B, Abundance. C, Shannon diversity index. D, Pielou evenness index.

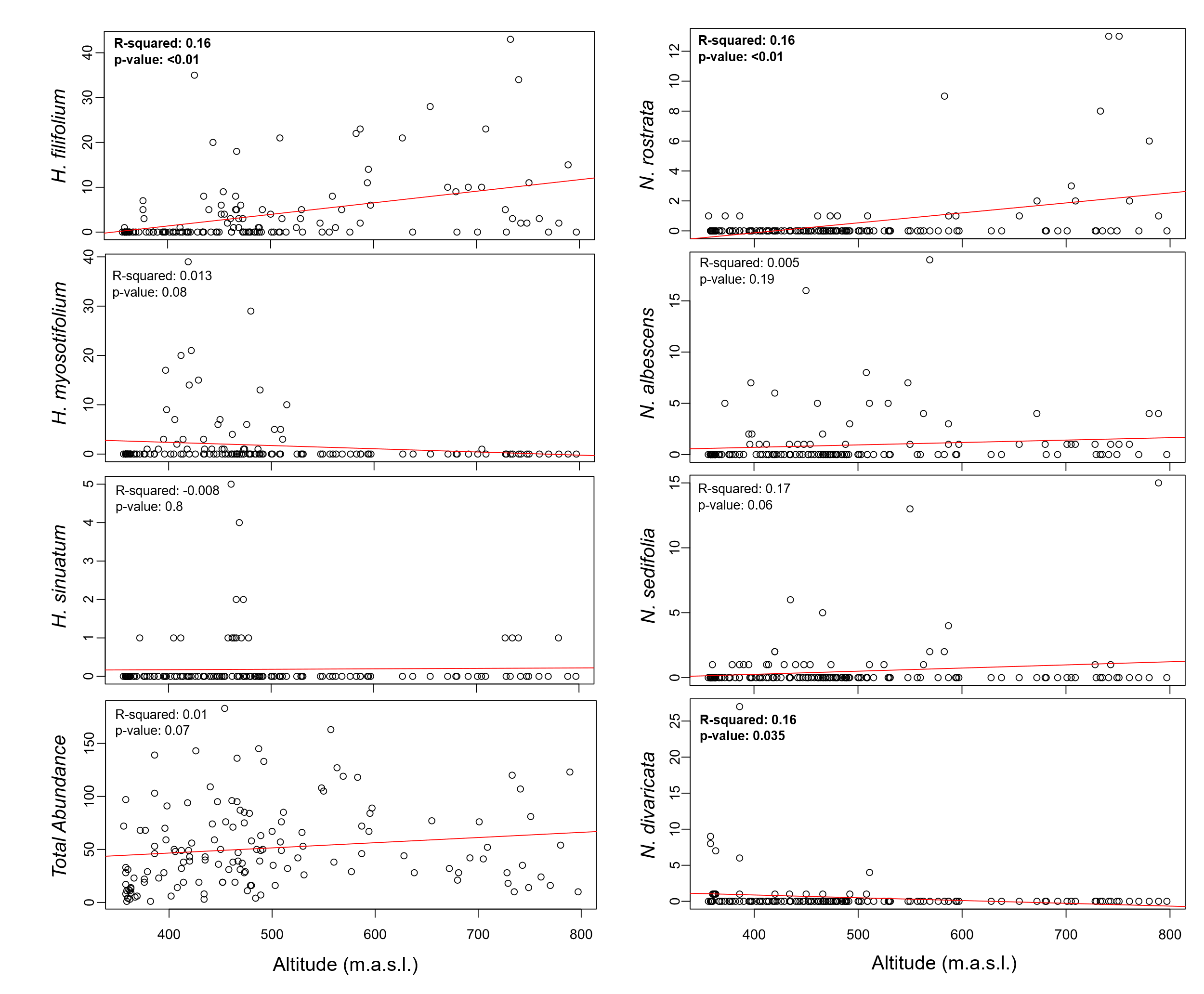

Figure S6. Linear regression between congeners abundance and elevation. Bold letters reflect significative relations.

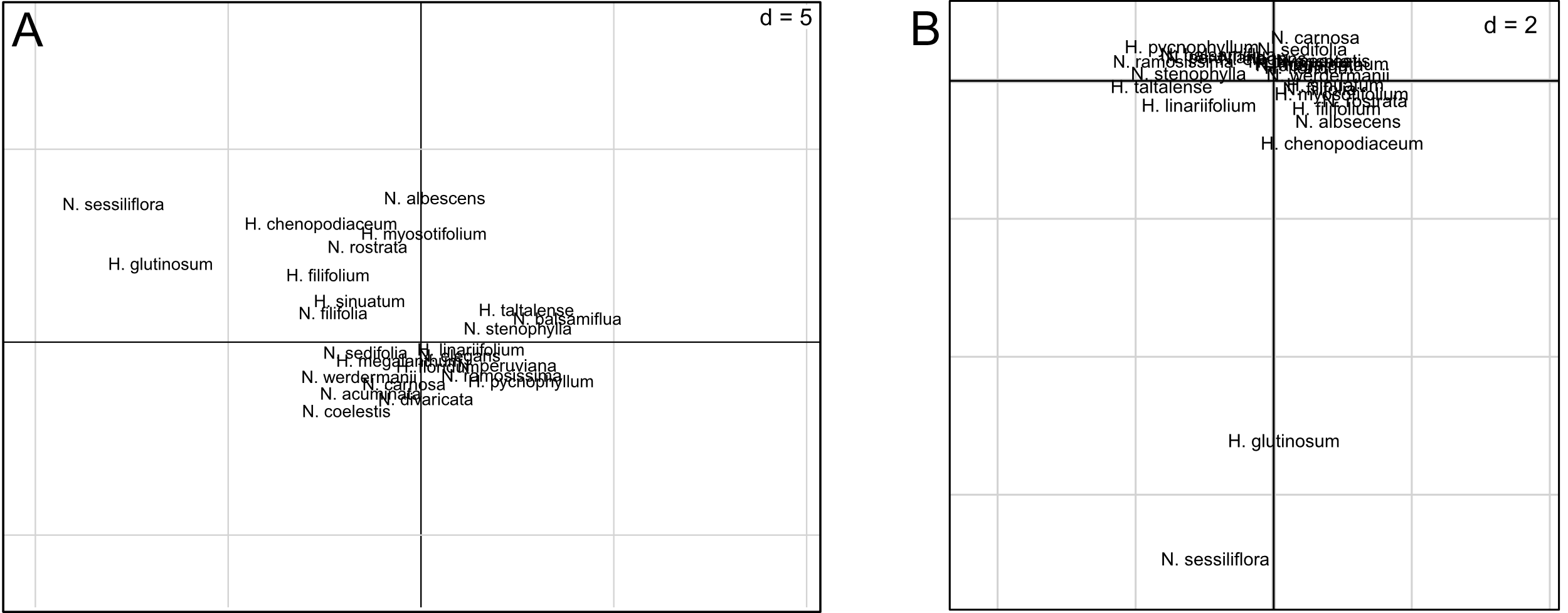

Figure S7. Placement of 15 and 10 species of *Nolana* and *Cochranea*, respectively, in the climatic multivariate space.

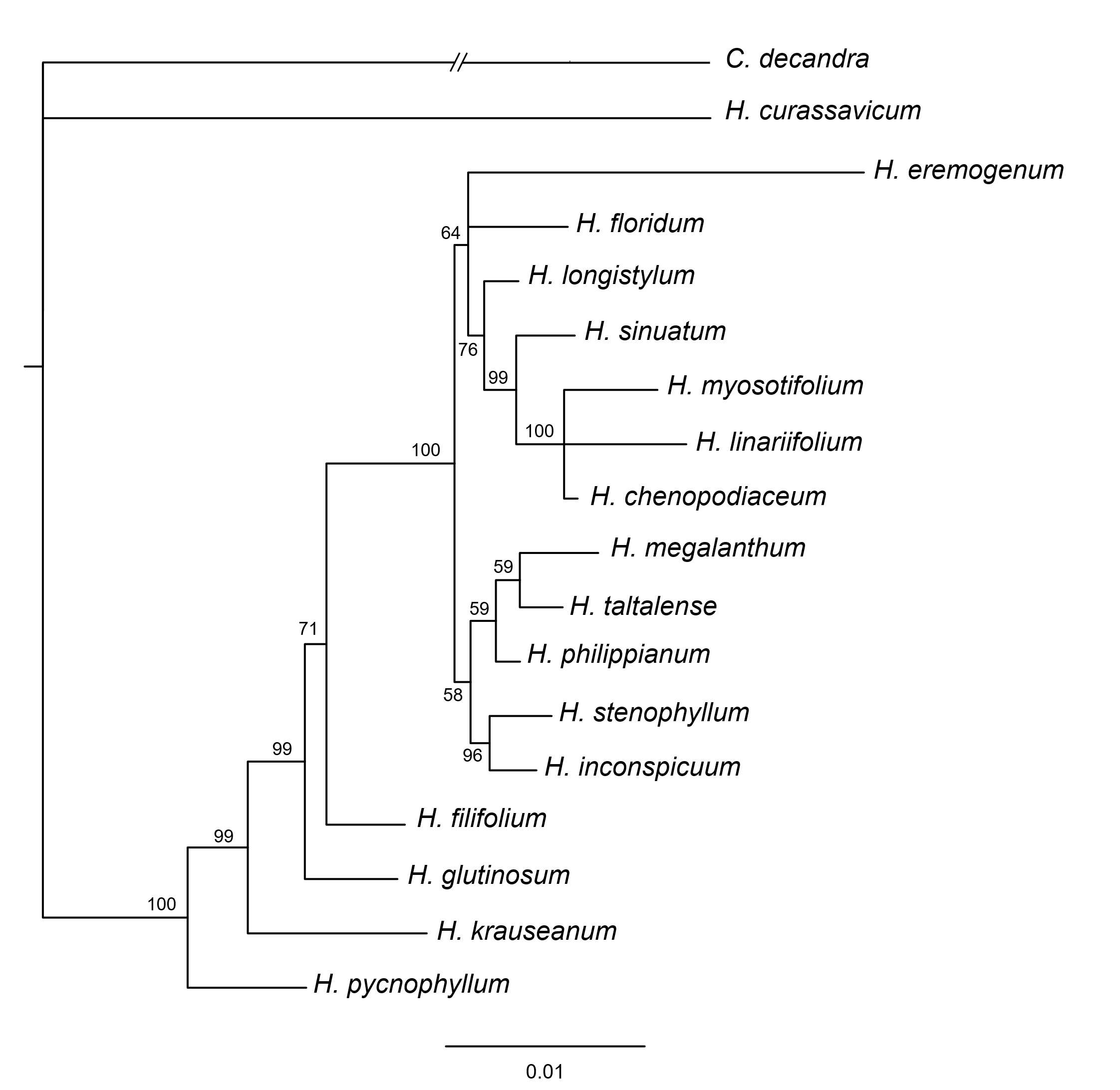

Figure S8. Phylogenetic relations of *Cochranea* based on one nuclear locus and four plastid loci. Bayesian posterior probabilities (PP) are shown next to the branches.

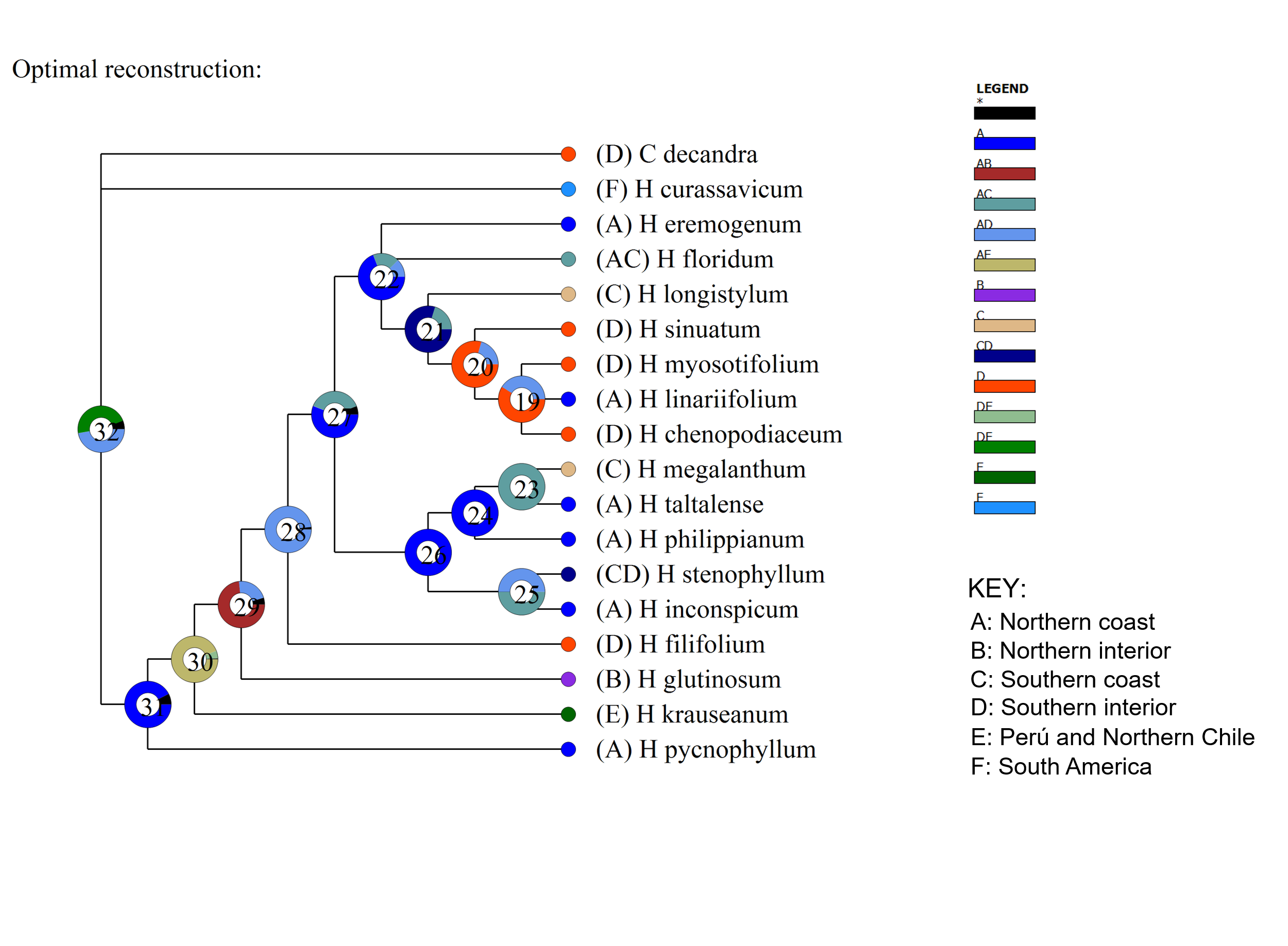

Figure S9. Ancestral area reconstruction of *Cochranea* using S-DIVA.

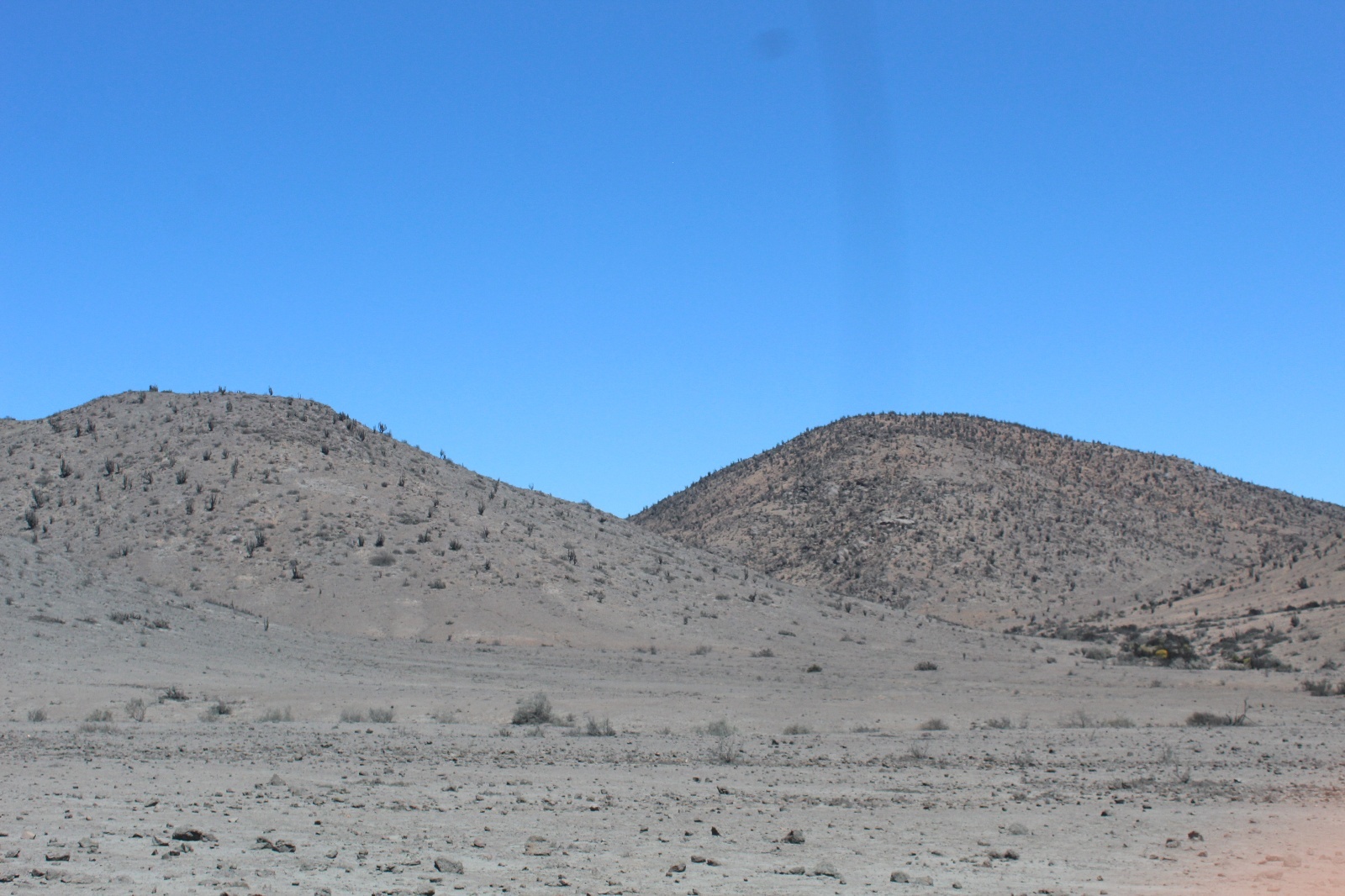

Figure S10. Decrease in vegetation density at the foot of the hills.

Figure S11. Presence data of *Nolana* and *Cochranea* species employed in climatic niche analyses. Red points, Northern interior. Yellow points, Northern coast; Blue points, Southern coast; Green points, Southern interior.

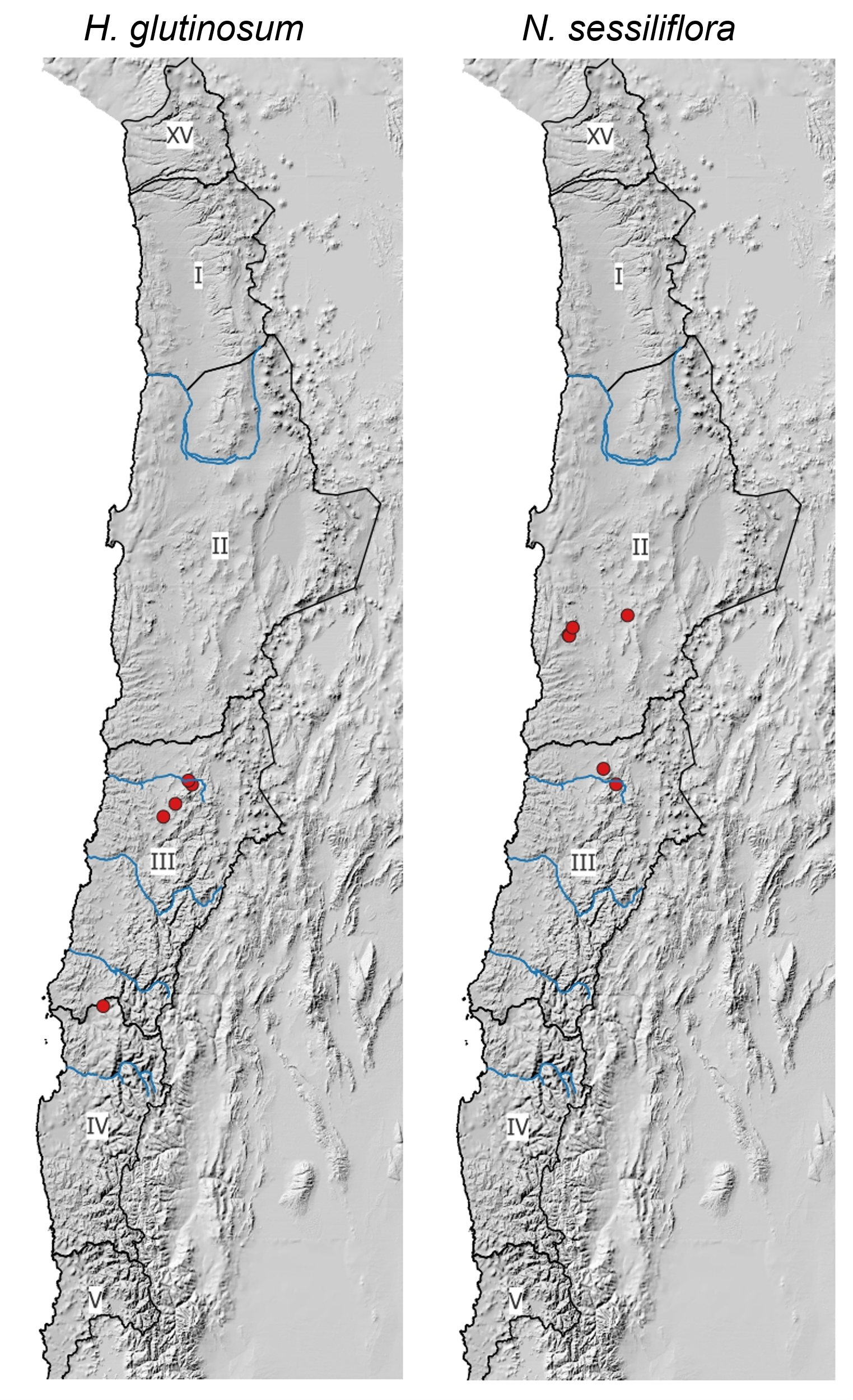

Figure S11. Continued.

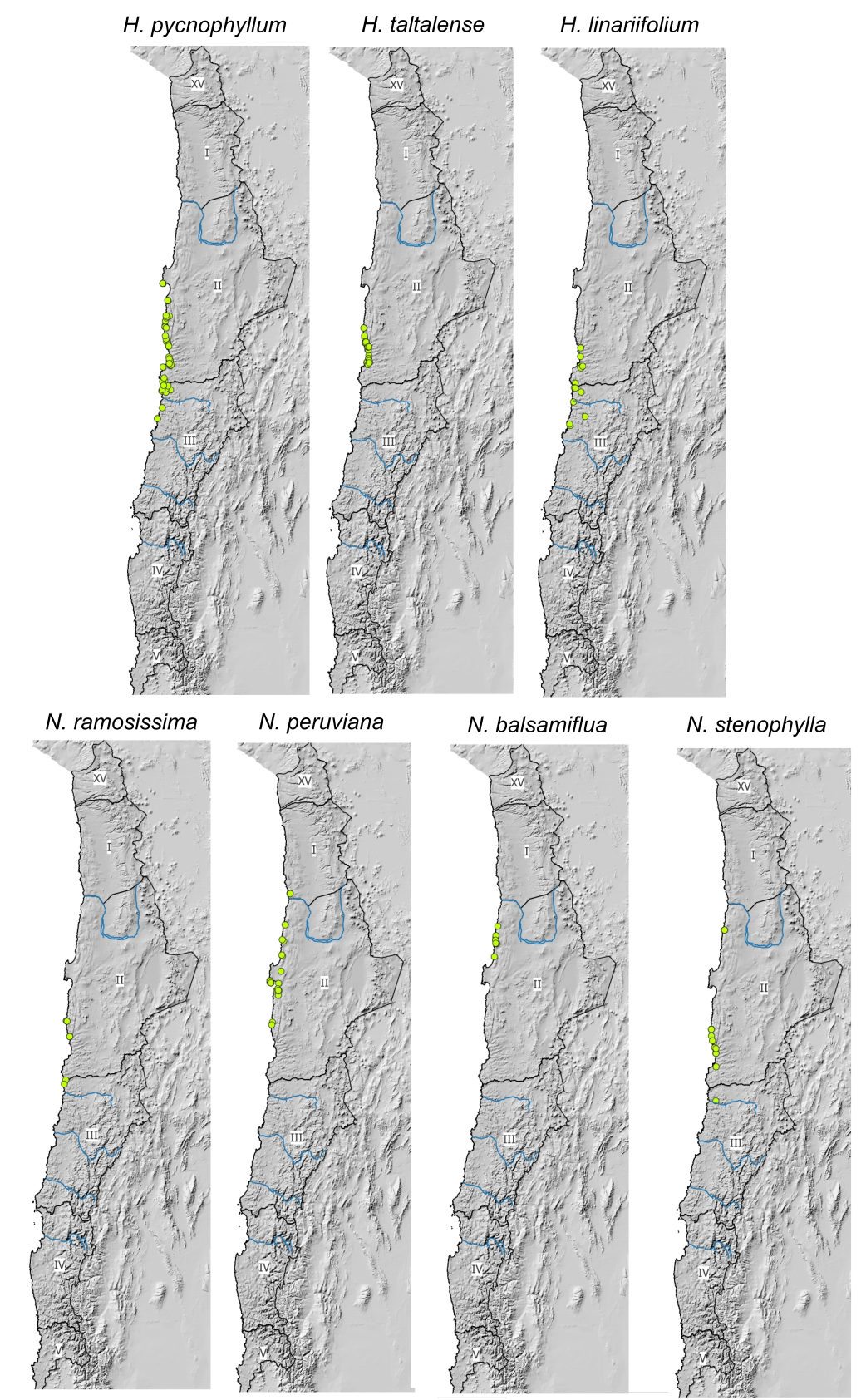

Figure S11. Continued.

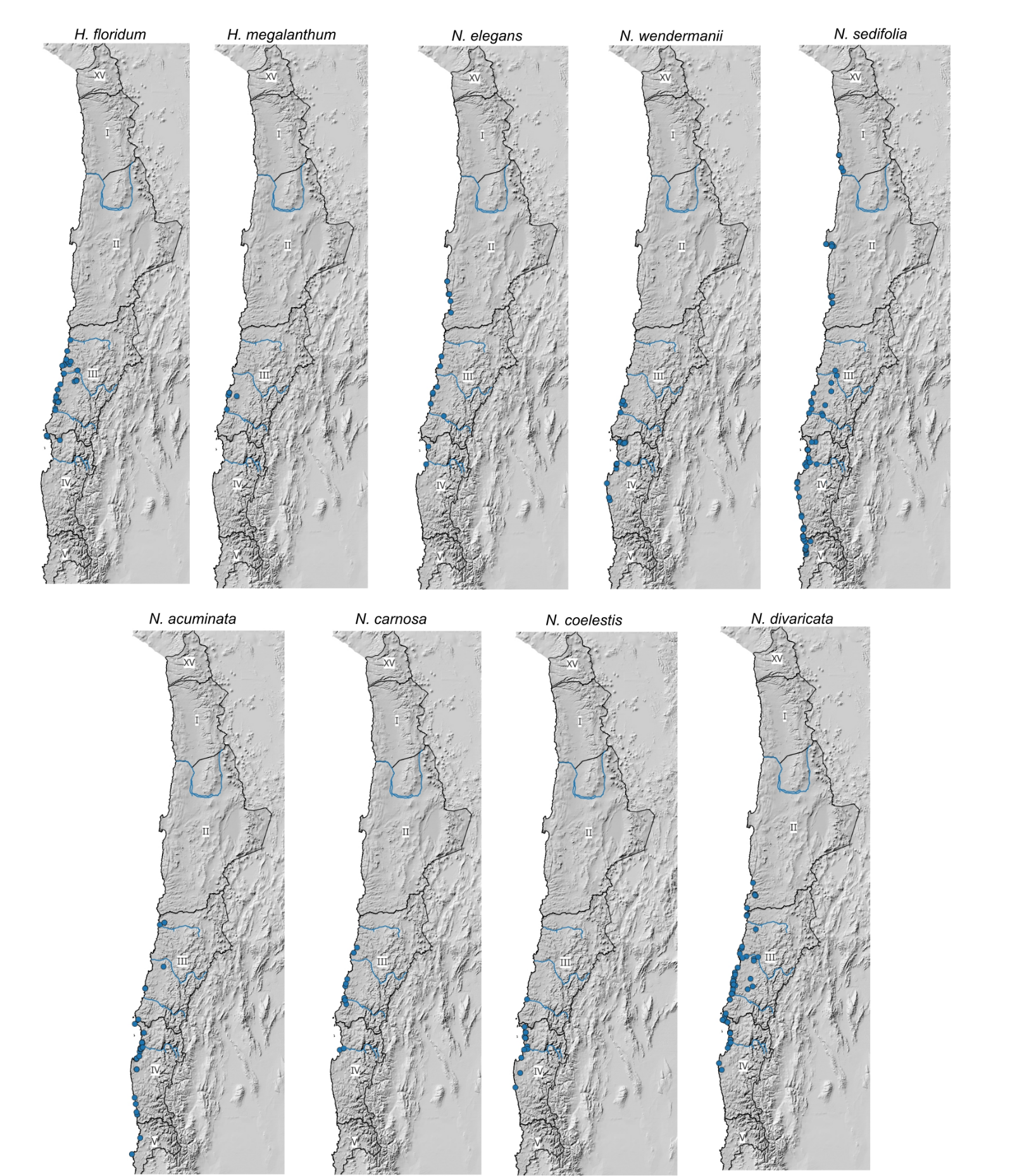

Figure S11. Continued.

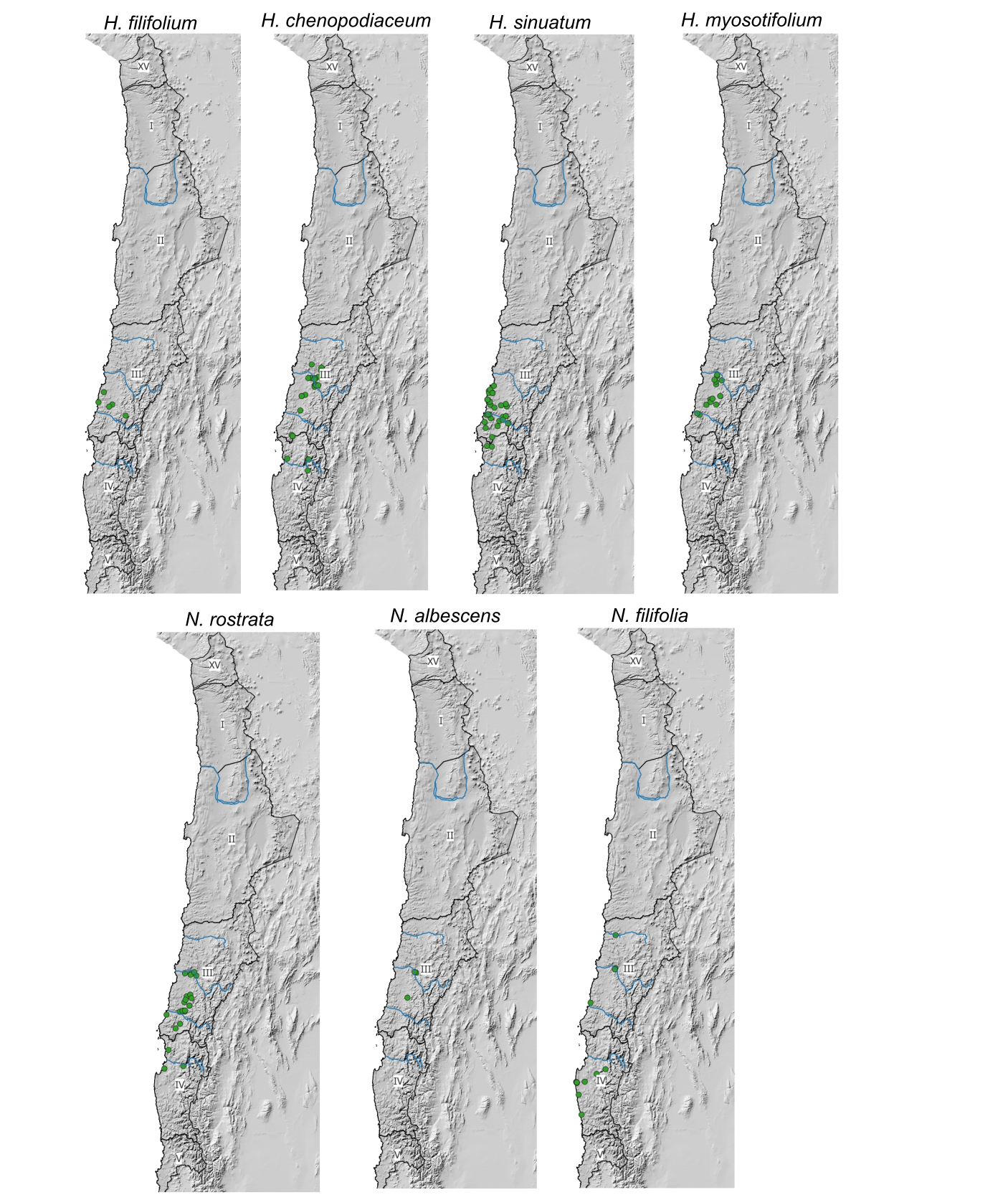

Appendix S1. R2 values of nMDS analysis representing the relationships between species and ordination axes.

***VECTORS

NMDS1	NMDS2	r2 Pr(>r)

Adesmia argentea	0.09755 -0.99523 0.2212 0.001 ***
Adesmia eremophila	0.31788 -0.94813 0.0361 0.068 .
Adiantum chilense	-0.52216 0.85285 0.0100 0.495
Argylia radiata	0.98218 -0.18793 0.0153 0.261
Atriplex deserticola	-0.98196 0.18907 0.0469 0.044 *
Balbisia peduncularis	-0.86285 -0.50546 0.0560 0.021 *
Balsamocarpon brevifolium	-0.94656 0.32252 0.0780 0.016 *
Erythrostemon angulatus	0.32214 -0.94669 0.0942 0.004 **
Centaurea chilensis	-0.95253 0.30445 0.0530 0.032 *
Cheilanthes mollis	-0.75414 -0.65672 0.0184 0.194
Cordia decandra	-0.16775 -0.98583 0.0811 0.008 **
Copiapoa echinoides	-0.63807 0.76998 0.1182 0.002 **
Cumulopuntia sphaerica	-0.49746 -0.86749 0.0218 0.224
Encelia canescens	0.46895 -0.88322 0.2487 0.001 ***
Ephedra gracilis	-0.26462 -0.96435 0.0609 0.016 *
Errazurizia multifoliolata	0.00820 -0.99997 0.0335 0.091 .
Eulychnia acida	-0.81782 0.57548 0.3885 0.001 ***
Fagonia chilensis	0.94955 -0.31361 0.0228 0.173
Flourensia thurifera	-0.97473 -0.22340 0.0295 0.090 .
Frankenia chilensis	-0.48623 -0.87383 0.0097 0.486
Galenia pubescens	0.04581 -0.99895 0.0117 0.408
Heliotropium filifolium	-0.80494 0.59335 0.1283 0.002 **
Heliotropium myosotifolium	-0.37249 -0.92804 0.0738 0.012 *
Heliotropium sinuatum	-0.31360 -0.94956 0.0612 0.019 *
Krameria cistoidea	-0.84298 -0.53795 0.0644 0.007 **
Lycium bridgesii	-0.79890 -0.60147 0.0477 0.028 *
Lycium minutifolium	-0.21197 -0.97728 0.0320 0.096 .
Miqueliopuntia miquelii	-0.81486 0.57965 0.1827 0.001 ***
Nicotiana glauca	-0.31151 -0.95024 0.0457 0.023 *
Nolana albescens	-0.64155 -0.76708 0.1326 0.001 ***
Nolana divaricata	0.90093 0.43395 0.0425 0.038 *
Nolana rostrata	-0.87059 0.49201 0.0793 0.007 **
Nolana sedifolia	-0.99407 0.10875 0.0246 0.164
Ophryosporus triangularis	-0.78260 -0.62253 0.0160 0.302
Oxalis gigantea	0.34353 -0.93914 0.0049 0.693
Pleurophora pungens	0.31862 -0.94788 0.0560 0.015 *
Polyachyrus poeppigii	-0.99054 0.13723 0.0296 0.073 .
Senecio myriophyllus	-0.34645 -0.93807 0.0885 0.005 **
Senna cumingii	0.10873 -0.99407 0.1711 0.002 **
Skytanthus acutus	0.84632 0.53268 0.2979 0.001 ***
Solanum remyanum	-0.53163 -0.84698 0.0704 0.010 **
Spinoliva ilicifolia	-0.24962 -0.96834 0.0502 0.037 *
Tetragonia angustifolia	-0.97207 -0.23470 0.0395 0.040 *
Tillandsia landbeckii	-0.89226 0.45152 0.0825 0.006 **
Tropaeolum tricolor	-0.75907 0.65101 0.0102 0.458
Tweedia birrostrata	-0.06968 -0.99757 0.0288 0.097 .

---

Signif. codes: 0 ‘***’ 0.001 ‘**’ 0.01 ‘*’ 0.05 ‘.’ 0.1 ‘ ’ 1

Permutation: free

Number of permutations: 999

Appendix S2. Results of the Multilevel pattern analysis.

Multilevel pattern analysis

Association function: IndVal.g
 Significance level (alpha): 0.05
 Total number of species: 46
 Selected number of species: 16
 Number of species associated to 1 group: 16
 Number of species associated to 2 groups: 0
 Number of species associated to 3 groups: 0
 Number of species associated to 4 groups: 0
 Number of species associated to 5 groups: 0
 Number of species associated to 6 groups: 0
 Number of species associated to 7 groups: 0

List of species associated to each combination:

Group 3 #sps. 1
 A	B	stat	p.value

Heliotropium myosotifolium	0.7831 0.9412	0.859	0.001 ***

Group 4 #sps. 2
 A	B	stat	p.value

Encelia canescens	0.5552 1.0000	0.745	0.001 ***
Senna cumingii	0.6103 0.4286	0.511	0.017 *

Group 8 #sps. 5
 A	B	stat	p.value

Cumulopuntia sphaerica	0.4943 1.0000	0.703	0.001 ***
Nolana albescens	0.4356 0.7500	0.572	0.013 *
Krameria cistoidea	0.4347 0.7500	0.571	0.011 *
Erythrostemon angulatus	0.4724 0.6250	0.543	0.021 *
Tetragonia angustifolia	0.4664 0.5000	0.483	0.043 *

Group 9 #sps. 6
 A	B	stat	p.value

Miqueliopuntia miquelii	0.5271 0.9730	0.716	0.001 ***
Heliotropium filifolium	0.6297 0.7838	0.703	0.003 **
Eulychnia acida	0.4379 0.9730	0.653	0.001 ***
Nolana rostrata	0.8651 0.3784	0.572	0.014 *
Tillandsia landbeckii	1.0000 0.2432	0.493	0.027 *
Copiapoa echinoides	0.5180 0.4324	0.473	0.040 *

Group 10 #sps. 1
 A	B	stat	p.value

Skytanthus acutus	0.8998 0.5833	0.725	0.001 ***

---

Signif. codes: 0 ‘***’ 0.001 ‘**’ 0.01 ‘*’ 0.05 ‘.’ 0.1 ‘ ’ 1

Appendix S3. Results of the Discriminant analysis, with 4 and 3 climatic zones. NI, Northern interior; NC, Northern coast; SI, Southern Interior; SC, Southern coast.

Confusion Matrix and Statistics

NI	NC	SI	SC
 NI	10	0	1	0
 NC	0	203	1	16
 SI	0	1	84	38
 SC	0	29	25	275

Overall Statistics

Accuracy: 0.8375
 95% CI: (0.8076, 0.8644)
 No Information Rate: 0.4817
 P-Value [Acc > NIR]: < 2.2e-16

Kappa: 0.7415

Mcnemar's Test P-Value: NA

Confusion Matrix and Statistics

NI	NC	SI+SC
 NI	10	0	1
 NC	0	197	7
 SI+SC	0	35	433

Overall Statistics

Accuracy: 0.937
 95% CI: (0.9161, 0.9541)
 No Information Rate: 0.6457
 P-Value [Acc > NIR]: < 2.2e-16

Kappa: 0.8619

Mcnemar's Test P-Value: NA
